# Anticodon Engineered Transfer RNAs (ACE-tRNAs) are a Platform Technology for Suppressing Nonsense Mutations

**DOI:** 10.1101/2024.06.06.597760

**Authors:** Wooree Ko, Joseph J. Porter, Sacha Spelier, Tyler Couch, Isabelle van der Windt, Priyanka Bhatt, Kevin Coote, Martin Mense, Jeffrey M. Beekman, John D. Lueck

## Abstract

Nonsense mutations arise from single nucleotide substitutions that result in premature termination codons (PTCs). PTCs result in little to no full-length protein production and loss of mRNA expression through the nonsense-mediated mRNA decay (NMD) pathway. We demonstrate that anticodon engineered (ACE-) tRNAs efficiently suppress the most prevalent cystic fibrosis (CF) causing PTCs, promoting significant rescue of endogenous cystic fibrosis transmembrane conductance regulator (CFTR) transcript abundance and channel function in different model systems. We demonstrate that our best-performing ACE-tRNA, that decodes all UGA PTCs to a leucine amino acid, markedly rescues CFTR channel function from the most prevalent CF causing PTCs that arise from non-leucine encoding codons. Using this single ACE-tRNA variant, we demonstrate significant rescue of CFTR channel function in an immortalized airway cell line and two different primary CF patient-derived intestinal cell models with CFTR nonsense mutations. Thus, ACE-tRNAs have promise as a platform therapeutic for CF and other nonsense-associated diseases.

## INTRODUCTION

Nonsense mutations arise from a single nucleotide substitution of an amino acid encoding codon that results in one of three in-frame termination codons (TGA, TAG and TAA), so-called premature termination codons (PTCs)^1^. PTCs result in truncated proteins that often have complete loss of function and are associated with severe disease phenotypes^2^. Further, mRNAs that contain PTCs are rapidly degraded through activation of a quality control surveillance pathway called nonsense-mediated mRNA decay (NMD)^3^. While each nonsense-associated disease is generally considered rare, together ∼11% of all known mutations that cause disease are PTC mutations, affecting hundreds of millions of individuals worldwide^4^.

Cystic fibrosis (CF) is a relatively common progressive disorder that arises from ∼2,000 mutations that result in reduced or complete loss of cystic fibrosis transmembrane conductance regulator (CFTR) channel function^5–7^. CFTR is localized at the apical membrane of epithelial cells of several organs, including lung and pancreas, and conducts chloride and bicarbonate which are crucial for maintaining hydration and pH of the apical extracellular space^8,9^. CF mutations are categorized into six different classes based on how they influence CFTR biology^10,11^. Through a process called theratyping^12^, several CF mutations that result in maturation defect (class II), gating defect (class III), conductance defect (class IV), reduced quantity (class V) and reduced stability (class VI) have been found to be responsive to CFTR modulator mono- or combinatorial pharmacotherapies^10^. A requirement for rescue of CFTR function by CFTR modulators is the production of sufficient CFTR protein for the modulators to act on. For this reason, nonsense mutations (Class I), which affect ∼10% of individuals with CF, are not candidates for current modulator pharmacotherapies.

Several strategies are currently being explored for treatment of CF resulting from PTCs. Outside of gene complementation approaches, there has been considerable effort in developing drugs that promote PTC readthrough to restore full-length functional protein^4,13^. Aminoglycosides are the first class of readthrough agents identified^14^, which target the ribosomal decoding center and promote misincorporation of near-cognate aminoacyl-tRNAs at PTCs and generation of full-length protein^4,15^. Several readthrough compounds, also known as translational readthrough-inducing drugs (TRIDS), have been investigated for their therapeutic promise. While TRIDs have different modes of action on protein translation and translation termination processes, they all result in the misincorporation of amino acids at the PTCs to extend translation (reviewed in^13,16^). To date, no TRID has been approved for use in the clinic. This is not because of lack of PTC readthrough efficiency, but rather their low efficacy *in vivo*, toxicity or both (reviewed in ^16^). However, because TRIDs are generally agnostic to the gene harboring the nonsense mutation, they remain an attractive therapeutic approach for several or most diseases that result from PTCs.

Suppressor (sup-) tRNAs are a gene therapy PTC suppression approach that hold various advantages in comparison to small molecule based readthrough strategies. Sup-tRNAs are tRNA sequences with anticodons that have been purposefully engineered, so called anticodon engineered (ACE-) tRNAs, to recognize and efficiently suppress one of three stop codons (UAA, UAG or UGA) with their cognate amino acid, resulting in translation elongation at the PTC site and generation of full-length protein^17–22^. Important for their use therapeutically, ACE-tRNAs are a relatively small genetic cargo (∼76nt RNA and as small as 155bp transgene) that have been delivered as RNA, naked cDNA and with therapeutic viruses to target tissues^19,22–24^. We and others have demonstrated that ACE-tRNAs minimally suppress natural termination codons (NTCs), therefore mitigating the main concern of their cellular toxicity^20,22,25^. Further, delivery of ACE-tRNAs has been shown to promote efficient suppression of PTCs encoded in their native genomic context of *CFTR* in cultured human cells^24,26^ and *Idua* (α-L-iduronidase) in a mouse model for Hurler syndrome^22^, resulting in significant restoration of transcript abundance through inhibition of NMD and expression of full-length functional protein.

Here, we investigate the ability of ACE-tRNAs to rescue endogenous CFTR function in cultured human cell models of the most prevalent CFTR nonsense mutations. Further, we assess combinatorial treatment of CFTR modulators and NMD inhibition to enhance ACE-tRNA rescue of transcript abundance and channel activity. While our results indicate that CFTR modulators enhance ACE-tRNA rescued CFTR function in a PTC dependent manner, inhibition of NMD only provided a modest boost in CFTR channel activity. Next, we sought to determine the ability of one ACE-tRNA sequence to rescue CFTR function from a multitude of PTCs, thus potentially reducing the necessity to advance several ACE-tRNA sequences as CF therapeutics. In three different CF cell models, we demonstrate that a single leucine ACE-tRNA sequence effectively restores CFTR function from endogenous transcripts harboring four prevalent *CFTR* PTC mutations (G542X, R553X, R1162X and W1282X), accounting for >70% of all individuals with nonsense-associated CF. Further, we show for select PTCs, combinatorial treatment with CFTR modulators and NMD inhibition enhances rescued CFTR channel function. Overall, we report the promise of ACE-tRNAs as a platform therapeutic, where one therapeutic entity can be applied to several nonsense mutations that cause CF and other diseases.

## RESULTS

### ACE-tRNA-mediated CFTR PTC suppression was enhanced by hCAR expression in cultured human airway cells

To evaluate the suppression activity of our ACE-tRNA technology at nonsense mutations, we utilized the CFF-16HBEge (16HBEge) cell model system, which was generated by CRISPR/Cas9 gene editing of parental 16HBE14o- (WT) cells^27^. 16HBEge cell lines harboring *CFTR* nonsense mutations G542X (2.5%), R1162X (0.4%) or W1282X (1.2%) were selected for study with ACE-tRNAs because of their high allelic frequency in CF PTC patients^7^. Additionally, of the 23 different nucleotide substitutions that result in nonsense mutations, the most frequent (21%) arise from arginine codons (CGA→TGA)^1,28^. To generate WT CFTR channels we chose ACE-tRNA^Gly^ and ACE-tRNA^Arg^ to suppress G542X and R1162X PTCs, respectively. Because UGA suppressing ACE-tRNA^Trp^ is not highly active^21^, efficient suppression of W1282X with the cognate tryptophan is not feasible. However, we have previously demonstrated that leucine incorporation at the CFTR-W1282 (W1282L) by our highly active ACE-tRNA^Leu^ restores significant CFTR function^26^. We therefore adopted this approach in the current study.

To assess the ability of ACE-tRNAs to suppress CFTR PTCs in several cell types, we opted for viral delivery of ACE-tRNAs. We first assessed the adenovirus (Ad) serotype 5 transduction efficiency in WT 16HBE14o-cells with and without exogenous expression of the human coxsackievirus and adenovirus receptor (hCAR) using flow cytometry (see methods for *piggyBac* generation of stable hCAR 16HBE cells). Exogenous expression of hCAR has shown to improve Ad transduction efficiency of airway cells and play a role in maintenance of tight junction formation^29^. All Ads in this study had either a green or red fluorescent protein (GFP or mCherry) encoded in the pAd vector under control of a CMV promoter. Cells were plated on cell culture treated dishes, transduced with an Ad encoding 4x cassettes of scrambled ACE-tRNA sequence (Ad-Scr) at increasing multiplicity of infection (MOI) of viral genomes (VG) per cell (MOI 1, 10, 100, 500), and analyzed 48hrs post transduction. Flow cytometry histograms of GFP fluorescence vs cell counts show that increasing Ad-Scr MOIs resulted in a concordant increase in GFP positive populations of both WT (Figure 1A) and WT + hCAR (Figure 1B) cells. While the percent cells transduced by Ad-Scr were significantly enhanced by exogenous hCAR expression at MOI 10 (37 ± 0.5% vs 48 ± 0.1%), 100 (80 ± 0.3% vs 88 ± 0.3%) and 500 (92 ± 0.2% vs 95 ± 1.2%), it was not overt (Figure 1C). However, the median fluorescence intensities (MFIs) were markedly increased with hCAR expression for MOI 100 (∼105%) and 500 (∼324%; Figure 1D), indicating that exogenous hCAR expression increased Ad VG cellular uptake. Based on these results, we generated hCAR expressing G542X-, R1162X- and W1282X-16HBEge cell lines to determine if PTC suppression efficiency by Ad-ACE-tRNAs could be further enhanced. First, we compared transduction efficiency of cells with Ad-Scr for all cell lines, and also with Ad-ACE-tRNA^Gly^ (Ad-Gly), Ad-ACE-tRNA^Arg^ (Ad-Arg) and Ad-ACE-tRNA^Leu^ (Ad-Leu) for G542X-, R1162X- and W1282X-16HBEge cells with and without exogenous hCAR, respectively (Figure S1A-S1L). Importantly, Ad-Scr, Ad-Gly, Ad-Arg and Ad-Leu transduced 16HBEge cell lines exhibited similar results as Ad-Scr transduced WT and WT + hCAR cell lines, with increased GFP MFIs with increased Ad MOIs and with the addition of hCAR. Next, we wanted to determine if the number of ACE-tRNA expression cassettes per Ad genome influenced PTC suppression efficiency. We transduced R1162X- (Figure S1M) and R1162X-16HBEge + hCAR (Figure S1N) cells with increasing Ad-Arg MOIs (MOI 1, 10 100 and 500) harboring 1 copy (Ad-1xArg), 4 copies (Ad-4xArg) and 8 copies (Ad-8xArg). Our ACE-tRNA expression cassette design used here is similar to our previously published studies^26^, containing a 55bp 5’ upstream control element, followed by the ACE-tRNA coding sequence and a 3’ trailer containing a processing and transcription termination^21,26^ (∼155bps in total). Restoration of CFTR transcript abundance from NMD was used as a readout for PTC suppression efficiency. We previously demonstrated that rescue of CFTR transcript abundance closely follows CFTR channel functional rescue and general PTC suppression efficiency^26^. With Ad-1xArg, Ad-4xArg and Ad-8xArg, there was a clear dose-dependent rescue of CFTR transcript abundance in R1162X-16HBEge (Figure S1M) and R1162X-16HBEge + hCAR (Figure S1N) cells, with Ad-4xArg in + hCAR cells being the top performing.

**Figure 1.**
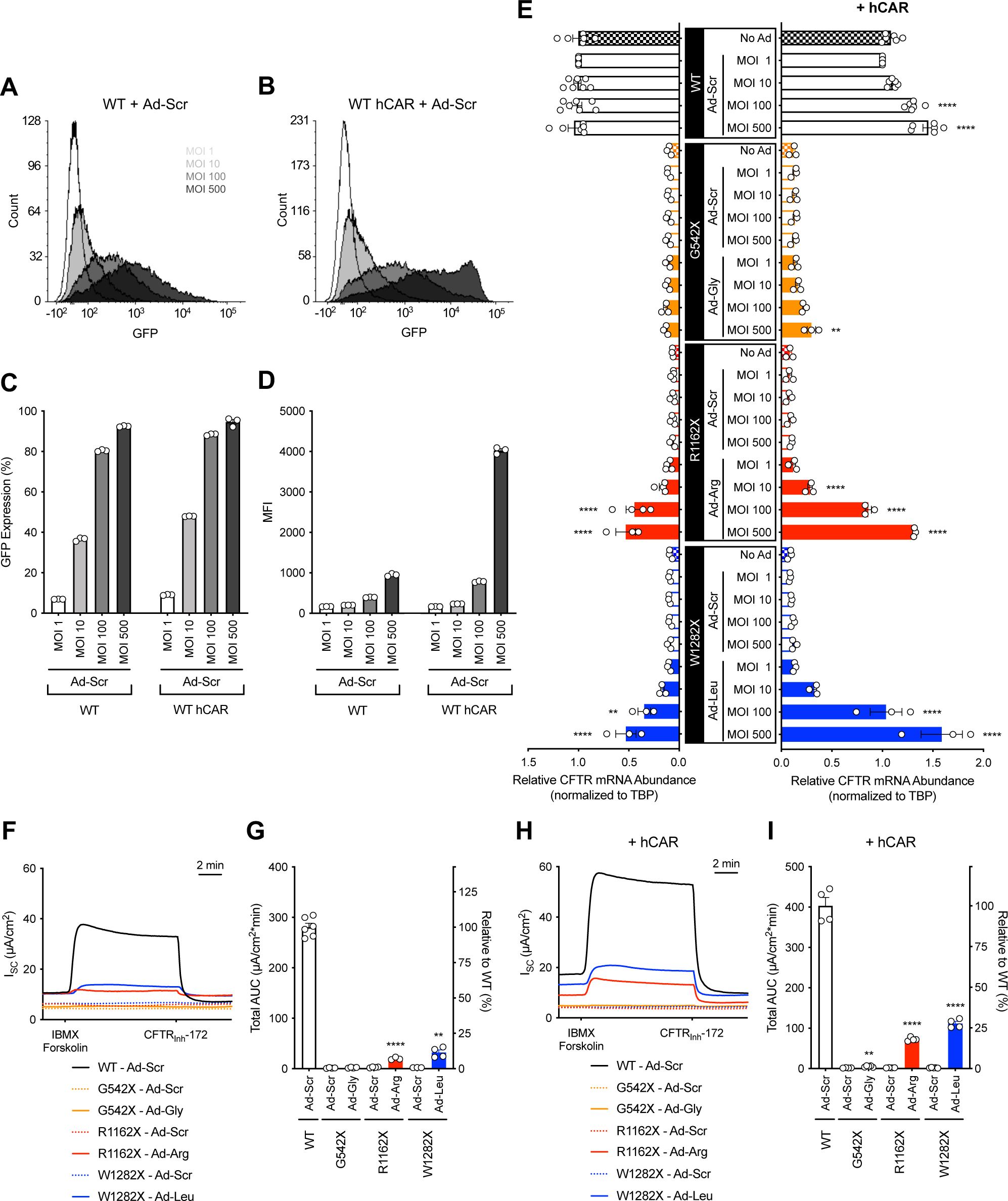
Stable expression of hCAR in 16HBE14o- and 16HBEge cell lines enhances adenovirus-mediated delivery of ACE-tRNAs. (**A-D**) Representative flow cytometry histograms of (**A**) WT or (**B**) hCAR expressing WT cells (WT + hCAR) transduced with an adenovirus (Ad) containing 4 copies of scrambled ACE-tRNA (Ad-Scr) and a cell transduction marker (GFP expression cassette). Quantification of (**C**) GFP positive cell population (*n* = 3) and (**D**) median fluorescence intensity (MFI) of that population (*n* = 3) was determined from flow cytometry data. Flow cytometry data for other cell lines are presented in **Figure S1A-L**. CFTR mRNA abundance (**E**) was normalized to TBP in WT (black), G542X-16HBEge (orange), R1162X-16HBEge (red) and W1282X-16HBEge (blue) cells (left) or the same set of 16HBE cells stably expressing hCAR (right). Cells were transduced at MOIs of 1, 10, 100 and 500 for each Ad-Scr (open bar), Ad-Gly (orange filled bar), Ad-Arg (red filled bar), or Ad-Leu (blue filled bar) (*n* = 3-7). CFTR channel function was determined by Ussing chamber recordings of (**F, G**) original or (**H, I**) + hCAR with Ads as indicated in the figure in response to addition of CFTR activators, forskolin (10 μM) and IBMX (100 μM), and then CFTR inhibitor, CFTR_Inh_-172 (20 μM). (**F, H**) Representative short-circuit Cl^−^ current (*I*_SC_) traces and (**G, I**) total area under the curve (AUC) quantification (*n* = 3-6). Data are presented as the mean ± SEM. Significance was determined by one-way ANOVA and Tukey’s post-hoc test, where ** *p* < 0.01 and **** *p* < 0.0001 vs. Ad-Scr MOI 1. Significance was determined by unpaired t-test for (**G**) and (**I**), where ** *p* < 0.01 and **** *p* < 0.0001.

We next sought to determine if our ACE-tRNA cassette design can be delivered to cells using adeno-associated virus (AAV) and efficiently suppress PTCs. We packaged 1x, 4x and 8x ACE-tRNA^Arg^ in AAV2 capsids (AAV-1xArg, AAV-4xArg and AAV-8xArg, respectively). Increasing MOIs (MOI 5,000, 10,000, 50,000 and 100,00) of AAV-1xArg, AAV-4xArg and AAV-8xArg were added to HEK293T cells that stably express a *piggyBac* transposon containing a Nanoluciferase (NLuc)-PTC luminescence protein and mScarlet-I fluorescent protein nonsense readthrough reporter system, which we call Stop-Go-Glow (SGG; Figure S2A). Of note, the NLuc-PTC has the context of hCFTR-W1282X with 30bps of human CFTR flanking sequence. After 48hrs post-transduction, the ability of AAV-1xArg, AAV-4xArg and AAV-8xArg to rescue NLuc expression was notable and dose-dependent, with MOI 100,000 yielding the highest NLuc signal in plate reader assays (Figure S2B). As expected, the control AAV-ACE-tRNA^Scr^ did not rescue NLuc expression at any MOI. Similar to our results with Ad, both the AAV-4xArg and AAV-8xArg gave similar dose-dependent rescue of NLuc expression. Using mScarlet-I expression as an indication of AAV2 transduction efficiency in flow cytometry assays, we found that increasing MOIs of AAV-1xArg, AAV-4xArg and AAV-8xArg resulted in increased transduction efficiency, with AAV-4xArg and AAV-8xArg, which outperformed AAV-1xArg, having similar dose-dependent transduction efficiencies (Figure S2C), mirroring results obtained with Ad. These results indicate that up to 8x ACE-tRNA expression cassettes can effectively be packaged in AAV capsids. Due to the cost of AAV production and the need to use high MOIs for transduction of cells in culture, we used Ad vectors expressing 4 copies of the ACE-tRNA expression cassettes for the remainder of studies, indicated as Ad-Scr, Ad-Gly, Ad-Arg and Ad-Leu for clarity.

When different ACE-tRNAs were tested in 16HBEge cell lines, transduction of R1162X-16HBEge cells with Ad-Arg and W1282X-16HBEge cells with Ad-Leu resulted in significantly increased CFTR transcript abundance at MOIs 100 and 500 (∼44% and ∼51% of WT for Ad-Arg MOIs 100 and 500, ∼34% and ∼51% of WT for Ad-Leu MOIs 100 and 500; Figure 1E, left) [Note: Rescue of transcript abundance comparisons were done to Ad-Scr transduced WT cells at matched MOIs]. The recovery of CFTR mRNA abundance is likely achieved through ACE-tRNA-mediated PTC suppression, completion of the pioneer round of translation and stabilization of the mRNA^30^. When hCAR expressing cells were transduced with Ads, significant recovery of CFTR transcript abundance was detected even with Ad-Gly in G542X-16HBEge + hCAR cells (∼21% of WT for Ad-Gly MOI 500) and near WT level of transcript abundance was rescued using MOIs 100 and 500 of Ad-Arg and Ad-Leu for R1162X-16HBEge + hCAR and W1282X-16HBEge + hCAR cells, respectively (∼66% and ∼91% of WT for Ad-Arg MOI 100 and 500, ∼80% and ∼109% of WT for Ad-Leu MOI 100 and 500; Figure 1E, right). Although MOI 500 of Ad vectors elicited the highest PTC suppression efficiency, we decided to use MOI 100 for the remainder of studies unless otherwise noted, as it supported ample PTC suppression and reduced the overall amount of Ad needed for the completion of the study. Together, these results indicate that Ad-mediated delivery of ACE-tRNAs inhibits NMD through efficient suppression of CFTR PTCs at three different sites in a dose-dependent manner, which is further enhanced by exogenous hCAR expression.

### Ad-ACE-tRNAs significantly rescued endogenous CFTR channel function in cultured human airway cells

Next, we determined whether Ad-ACE-tRNA-induced PTC suppression and significant rescue of transcript abundance translates to recovery of endogenous CFTR channel function. To accomplish this, we transduced 16HBEge cells with Ad-ACE-tRNA (MOI 100) 72hrs after seeding on permeable supports (Transwells) and performed Ussing chamber assays to measure forskolin/IBMX induced and CFTR_Inh_-172 inhibited transepithelial CFTR short-circuit current (*I*_sc_) at day 5 (reported as total area under the curve; AUC). Representative *I*_sc_ traces from G542X-16HBEge cells show that Ad-Gly was ineffective at CFTR functional rescue, whereas Ad-Arg and Ad-Leu transduction of R1162X-16HBEge and W1282X-16HBEge cells restored some CFTR activity (Figure 1F). Average AUC quantification is plotted in Figure 1G, with Ad-Arg and Ad-Leu transduction rescuing ∼7% and 15% of WT CFTR activity, respectively. [Note: Rescue of CFTR function comparisons were done to Ad-Scr transduced WT cells.] The disparity in the degree of functional CFTR rescue is the aggregate of differences in ACE-tRNA suppression “strength” (Arg>Leu>>Gly)^31^ and the impact of nucleotide context on PTC suppression efficiency^32,33^. In hCAR expressing cell lines, all three ACE-tRNAs significantly restored CFTR channel function, ∼1% with Ad-Gly, ∼18% with Ad-Arg and ∼28% with Ad-Leu of WT (Figure 1H and 1I). However, even with exogenous hCAR expression, the magnitude of CFTR channel rescue on Transwells was not consistent with that of rescued CFTR transcript abundance (Figure 1E). Using flow cytometry analysis of GFP fluorescence on cells following Ussing chamber measurements in paired experiments, we found that Ad transduction efficiencies for 16HBE (∼20%) and 16HBE + hCAR (∼34%) cell lines seeded on Transwells were markedly lower than observed with cells transduced on cell culture dishes, however they had comparable MFIs (Figure S1O-S1V). The expression of hCAR significantly boosted Ad transduction efficiency and was therefore used in all subsequent 16HBE experiments. While Ad-ACE-tRNA transduction efficiency was significantly impaired on Transwells, the amount of CFTR channel functional restoration tracked with delivery efficiencies of the highly active Ad-Arg and Ad-Leu. Together, these results demonstrate that when ACE-tRNA^Arg^ and ACE-tRNA^Leu^ are delivered with high efficiency, they can significantly restore levels of endogenous CFTR channel activity.

### CFTR modulators enhanced ACE-tRNA-mediated functional CFTR rescue in cultured human airway cells

To date, four CFTR-specific modulator treatments have been approved for the treatment of many CF patients with CFTR mutations in classes II-VI: ivacaftor (I), lumacaftor/ivacaftor (LI), tezacaftor/ivacaftor (TI) and elexacaftor/tezacaftor/ivacaftor (ETI)^34–37^. We hypothesized that ACE-tRNA-mediated recovery of CFTR channel expression would provide a target for CFTR modulators to further enhance CFTR channel function. To test this hypothesis, we delivered Ad-ACE-tRNAs to 16HBEge + hCAR cell lines seeded on Transwells and measured restoration of CFTR channel function, followed by transcript abundance in the presence of different combinations of CFTR modulators.

Despite significant differences in CFTR functional rescue, Ad-Arg and Ad-Leu transduction of R1162X-16HBEge (Figure 2A) and W1282X-16HBEge (Figure 2D) cells resulted in similar rescue of CFTR transcript abundance to 40-50% and 50-63% of WT, respectively, across all treatment conditions. Representative CFTR *I*_sc_ traces demonstrated a clear difference in the ability and extent of modulators to augment the function of ACE-tRNA-dependent rescued R1162X-CFTR (Figure 2B) and W1282X-CFTR (Figure 2E) channels. LI treatment resulted in the highest yet meager synergistic rescue of channel function with Ad-Arg (∼18% for LI of WT; Figure 2C). Different from R1162X-16HBEge cells, the best treatment combination for W1282X-16HBEge cells was Ad-Leu + ETI treatment, with the significant enhancement in CFTR channel function being evident from the representative *I*_sc_ traces (Figure 2E), with an average rescue of ∼50% of WT channel function (Figure 2F). The breakout analysis of the individual effects of Ad-ACE-tRNAs and each treatment (forskolin/ IBMX, modulators and CFTR_Inh_-172) on R1162X- and W1282X-16HBEge cells are presented in Figure S3A-D). It has been previously demonstrated that the CFTR-W1282L variant has WT CFTR trafficking and channel behavior that is enhanced by LI treatment^38^. CFTR-W1282L activity is potently enhanced by acute ivacaftor treatment following ET pre-treatment (Figure 2E and S3C), suggesting this variant is hypersensitive to ivacaftor potentiation. Importantly, unlike G418 treatment^27^, Ad-Arg or Ad-Leu transduction of WT 16HBEo-cells had no adverse effect on CFTR transcription abundance and channel function (Figure 2A-2F and S3A-S3D). This supports the growing body of evidence that ACE-tRNAs do not affect normal translation processes and are not toxic to cells.

**Figure 2.**
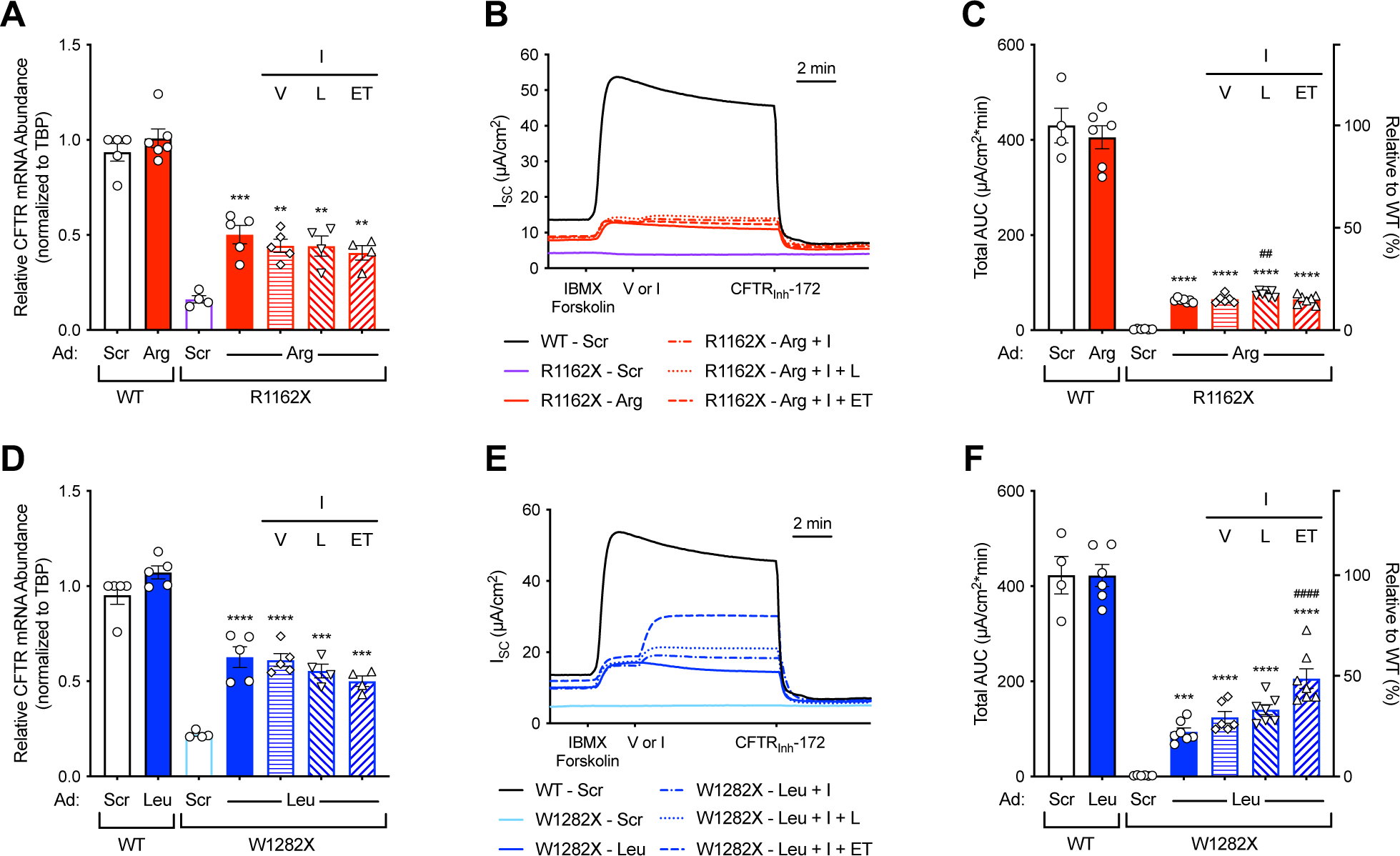
Combinatorial treatment of ACE-tRNAs and CFTR modulators sigenhances rescued CFTR channel function. (**A-C**) WT and R1162X-16HBEge cells (+ hCAR) were transduced with Ad-Scr or Ad-Arg (MOI 100), pretreated with vehicle (V), lumacaftor (L, 3 μM) or elexacaftor and tezacaftor (ET, 3 μM each) for 24 hours and then acutely treated with vehicle (V) or ivacaftor (I, 1 μM). (**A**) Relative CFTR mRNA abundance was measured in the same cell population after measurement of CFTR channel function by Ussing chamber (*n* = 4-6), (**B**) representative *I*_SC_ traces and (**C**) total AUC quantification (*n* = 4-7) of Ussing chamber recordings in response to sequential addition of forskolin (10 μM) and IBMX (100 μM), followed by either vehicle (V) or ivacaftor (I, 1 μM), and then by CFTR_Inh_-172 (20 μM). For (**A**) and (**C**), data are presented as mean ± SEM. (**D-F**) WT and W1282X cells (+ hCAR) were transduced with Ad-Scr or Ad-Leu (MOI 100) and treated identically to R1162X-16HBEge cells outlined above. (**D**) Relative CFTR mRNA abundance quantification was done following Ussing chamber measurements of CFTR channel function (*n* = 4-5), (**E**) representative *I*_SC_ traces and (**F**) total AUC quantification (*n* = 4-7) from cells treated identically to cells presented in (**C**). For (**A**), (**C**), (**D**), and (**E**), data are presented as the mean ± SEM. Significance was determined by unpaired t-test or one-way ANOVA and Tukey’s post-hoc test, where ** *p* < 0.01, *** *p* < 0.001 and **** *p* < 0.0001 vs. Ad-Scr, ^##^ *p* < 0.01 vs. Ad-Arg and ^####^ *p* < 0.0001 vs. Ad-Leu.

### ACE-tRNA-mediated rescue of CFTR function was enhanced by NMD inhibition

Next, we sought to determine if reduced transcript abundance through NMD was a limiting component of ACE-tRNA-dependent rescue of CFTR channel function. To test this idea, we inhibited NMD with SMG1 kinase inhibitor (SMG1i) in R1162X-16HBEge and W1282X-16HBEge cells seeded on Transwells that were transduced with Ad-Arg and Ad-Leu, respectively. SMG1i treatment increased CFTR transcript abundance similarly in R1162X-16HBEge cells transduced with Ad-Scr (∼10%; Figure 3A, white and gray filled purple bars) and Ad-Arg (∼13%; Figure 3A, white and gray filled red bars) with no effect of CFTR modulators (Figure 3A, hashed bars). Because SMG1i treatment had little impact on R1162X-CFTR transcript abundance both in the absence and presence of Ad-Arg, we were not surprised to find that R1162X-CFTR channel function was not significantly enhanced by its treatment (Figure 3B and 3D). Similarly, Galietta and colleagues reported that SMG1i treatment did not enhance functional rescue of CFTR in R1162X-16HBEge cells treated with readthrough molecules G418 or ELX-2^39^. Of note, restored R1162X-CFTR channel activity with combinatorial treatment of G418 or ELX-2, modulators and SMG1i was much less (<10% of WT) than what is achieved by Ad-Arg treatment alone in 16HBEge cells. Indeed, the most significant contribution of R1162X-CFTR functional recovery was Ad-Arg treatment (∼20% of WT), with combinatorial Ad-Arg, SMG1i and LI treatment recovering ∼29% of WT (Figure 3B, gray filled hashed red bar). Mechanistically, it is unclear why there is a disparity in enhancement of CFTR channel function by LI treatment of WT (∼41%) and generation of WT channels through Ad-Arg rescue of R1162X-CFTR channels (∼9%). The breakout analysis of the average contribution of all treatments on R1162X-CFTR channel function is displayed in Figures S3E and S3F, and representative CFTR transepithelial *I*_SC_ traces are presented in Figure 3D. The relationship between CFTR function and CFTR mRNA abundance for WT and R1162X-16HBEge cells are plotted in Figure 3C and highlights the disparity of LI enhancement of WT and Ad-Arg rescued R1162X-CFTR channels and NMD inhibition and rescue of R1162X-CFTR channel function.

**Figure 3.**
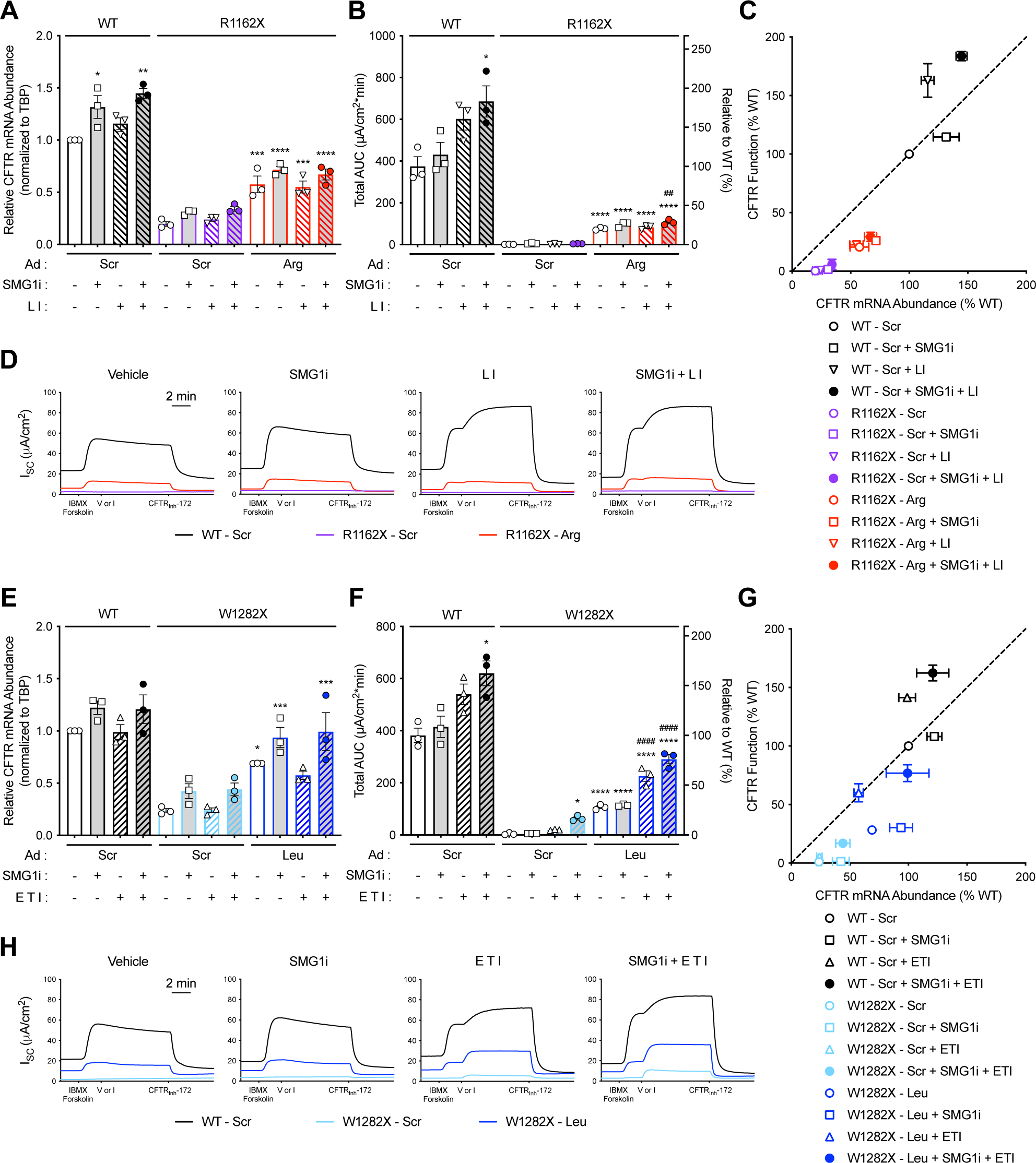
NMD inhibition and CFTR modulation significantly enhance ACE-tRNA-dependent rescue of CFTR channel function. (**A-D**) WT and R1162X-16HBEge (+ hCAR) were transduced with Ad-Scr (black for WT, purple for R1162X-16HBEge) or Ad-Arg (red) at an MOI of 100 and pretreated with either vehicle (V) or SMG1i (0.3 μM) for 48 hours and then with either vehicle (V) or lumacaftor (L, 3 μM) for 24 hours. Cells pretreated with L were acutely treated with ivacaftor (I, 1 μM). (**A**) Relative CFTR mRNA abundance was quantified in the same cell population after measurement of CFTR channel function by Ussing chamber (*n* = 3), (**B**) total AUC quantification (*n* = 3), (**C**) CFTR function in relation to CFTR mRNA abundance (*n* = 3) and (**D**) representative *I*_SC_ traces from Ussing chamber recordings in response to sequential addition of forskolin (10 μM) and IBMX (100 μM), followed by either vehicle (V) or ivacaftor (I, 1 μM), and then by CFTR_Inh_-172 (20 μM). (**E-H**) WT and W1282X-16HBEge cells (+ hCAR) were transduced with Ad-Scr (black for WT, light blue for W1282X-16HBEge) or Ad-Leu (blue) at an MOI of 100 and pretreated with either vehicle (V) or SMG1i (0.3 μM) for 48 hours and then with either vehicle (V) or ET (3 μM each) for 24 hours. Cells pretreated with ET were acutely treated with ivacaftor (I, 1 μM). (**E**) Relative CFTR mRNA abundance quantified in the same cell population after measurement of CFTR channel function by Ussing chamber (*n* = 3), (**F**) total AUC quantification (*n* = 3), (**G**) CFTR channel function in relation to CFTR mRNA abundance (*n* = 3) and (**H**) representative *I*_SC_ recordings from cells treated identical to those presented in (**D**). Data are presented as the mean ± SEM for (**A-C**, **E-G**). Breakout analysis for effect size of each treatment are shown in **Figure S3**. Significance was determined by one-way ANOVA and Tukey’s post-hoc test, where * *p* < 0.05, ** *p* < 0.01, *** *p* < 0.001 and **** *p* < 0.0001 vs. Ad-Scr, ^##^ *p* < 0.01 vs. Ad-Arg, and ^####^ *p* < 0.0001 vs. Ad-Leu.

Consistent with previous reports^27,39^, NMD of W1282X-CFTR transcripts were highly responsive to SMG1i treatment, recovering ∼19% of transcript abundance in W1282X-16HBEge cells transduced with Ad-Scr (∼24 to ∼43% of WT; Figure 3E; white and gray filled light blue bars) and enhanced Ad-Leu dependent rescue of transcript abundance by ∼25% (∼69 to ∼94% of WT; Figure 3A, white and gray filled blue bars). As observed with R1162X-CFTR, the combinatorial effect of Ad-Leu + SMG1i on restoration of W1282X-CFTR transcript abundance was additive and was not enhanced by addition of modulators (Figure 3E, hashed bars). SMG1i-dependent rescue of W1282X-CFTR transcript abundance did not translate to increased channel function alone or in combination with Ad-Leu (Figure 3F, white and gray filled blue bars). However, SMG1i + ETI combinatorial treatment markedly restored W1282X-CFTR channel function to ∼17% of WT (Figure 3F, hashed light blue bars). This is consistent with previous reports that increased transcript abundance results in increased truncated and/or full-length CFTR expression by spurious PTC readthrough for CFTR modulators to act on and restore functional CFTR channels^27,38–40^. Combinatorial treatment of W1282X-16HBEge cells with Ad-Leu + SMG1i + ETI resulted in an impressive rescue of W1282X-CFTR channel function to ∼76% of WT (Figure 3F, gray filled hashed blue bar). The breakout analysis of the average contribution of all treatments on W1282X-CFTR channel function is displayed in Figures S3G and S3H, and representative CFTR transepithelial *I*_SC_ traces are presented in Figure 3H. By plotting the relationship of W1282X-CFTR mRNA abundance and channel function, it becomes clear that rescue of transcript abundance by SMG1i provides only minimum benefit, with the majority of rescued channel activity resulting from combinatorial Ad-Leu and ETI treatment (Figure 3G).

We next asked whether treatment with CFTR modulators alone or in combination with SMG1i allows for a reduced Ad-ACE-tRNA MOI for significant CFTR channel functional rescue. As a corollary, we wanted to determine if increasing CFTR transcript abundance through NMD inhibition is disproportionally beneficial for rescue of CFTR channel function when ACE-tRNAs may be limiting with low MOIs. Ad-Gly, Ad-Arg and Ad-Leu at an MOI of 10 was used to transduce G542X-, R1162X- and W1282X-16HBEge cells, respectively. [Note: These studies were performed on cells plated on cell culture dishes.] For G542X- and W1282X-16HBEge cells, treatment with SMG1i alone significantly rescued CFTR transcript abundance (Figure S4A, white filled bars). In contrast, R1162X-16HBEge cells required the combination of Ad-Arg + SMG1i to significantly restore transcript abundance, again highlighting the positional effect of PTCs on rescue of CFTR transcripts from NMD by PTC readthrough or direct NMD inhibition^41^. Next, we performed a dose response of Ad-ACE-tRNAs (MOI 10, 50, 100 and 500) with CFTR modulators alone or CFTR modulators + SMG1i in R1162X- and W1282X-16HBEge cells seeded on Transwells for paired quantification of CFTR channel function and transcript abundance. In R1162X-16HBEge cells, there was little difference in rescued R1162X-CFTR transcript abundance at MOIs of 10 and 500, in the absence (Figure 4A, white hashed bars) or presence (Figure 4A, gray hashed bars) of SMG1i. This observation was similar in functional studies, with only a small (∼12%), albeit significant, increase in R1162X-CFTR channel function between Ad-Arg MOI 10 + LI (∼15% of WT) and Ad-Arg MOI 500 + LI +SMG1i (∼27% of WT; Figure 4B), supporting previous reports that very few ACE-tRNAs need to be delivered to provide significant PTC suppression activity^26^. As observed with R1162X-CFTR, Ad-Leu MOIs >50 did not support enhanced rescue of W1282X-CFTR transcript abundance (Figure 4C) or channel function (Figure 4D), in the absence (white hashed bars) or presence (gray hashed bars) of SMG1i. MOI 10 Ad-Leu + ETI + SMG1i was comparable to MOIs 100-500 Ad-Leu + ETI for rescue of both W1282X-CFTR transcript abundance (∼45 vs. ∼46% of WT; Figure 4C) and channel function (∼55 vs. ∼52% of WT; Figure 4D), suggesting that the abundance of CFTR transcript plays a small role in limiting ACE-tRNA-dependent rescue CFTR channel function. Further, there was not an outsized benefit of SMG1i treatment for rescue of CFTR transcript abundance or channel function at MOI 10 Ad-Leu and Ad-Arg compared to MOI 500. The breakout analysis of the impact of each treatment on CFTR channel function is displayed in Figures S4B-S4E, along with representative *I*_sc_ traces in Figures S4F and S4G. Taken together, results demonstrate that even at low MOIs, Ad-ACE-tRNAs alone efficiently rescue endogenous R1162X-CFTR and W1282X-CFTR channel function. Further, combinatorial ACE-tRNA treatment with CFTR modulators, and to a lesser extent NMD inhibition, may be viable options for responsive CF nonsense variants like W1282X.

**Figure 4.**
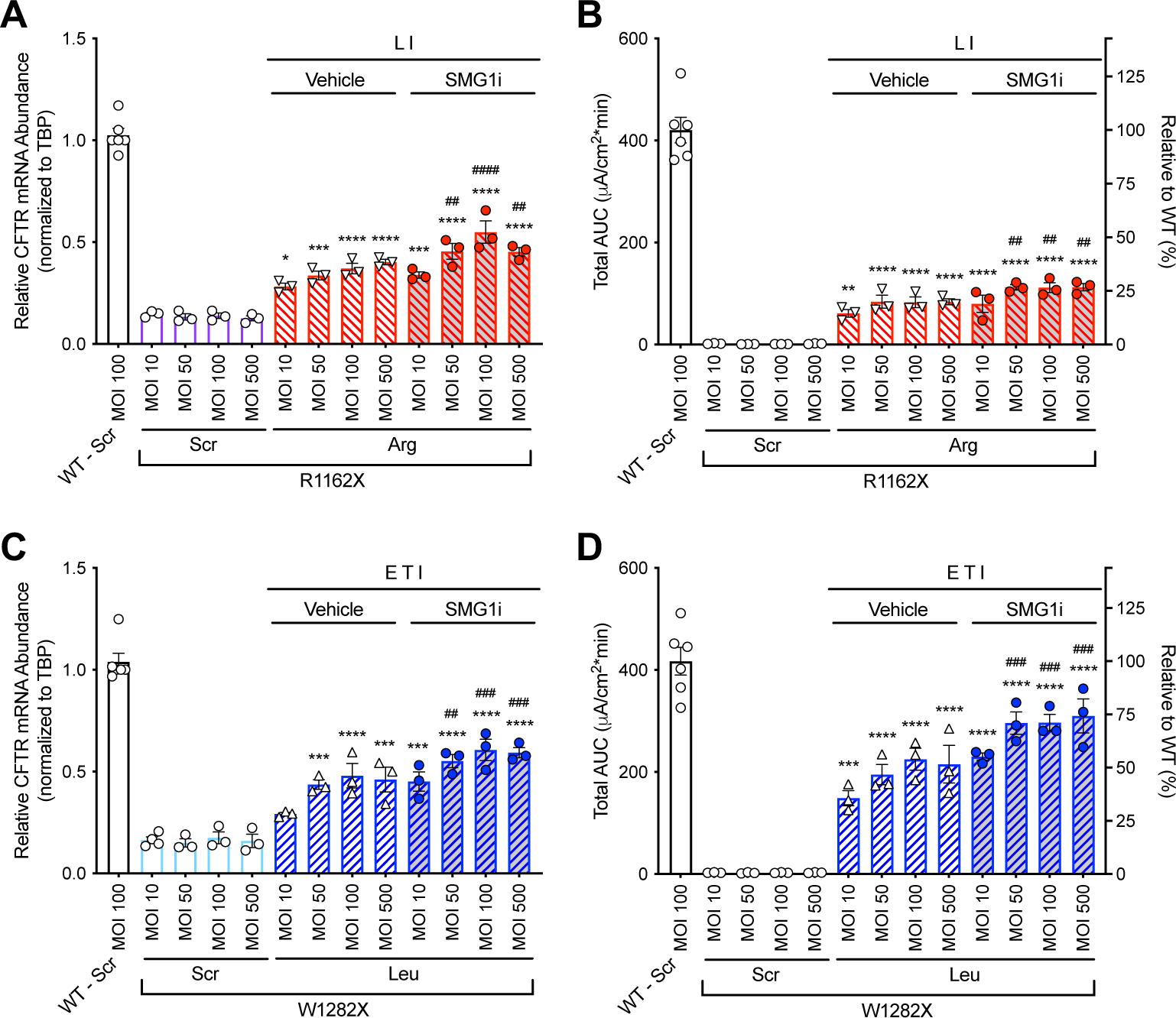
Use of CFTR modulators and SMG1i NMD inhibition significantly reduces the ACE-tRNA dosage required for significant CFTR functional rescue. (**A, B**) R1162X-16HBEge and (**C, D**) W1282X-16HBEge cells (+ hCAR) transduced with Ad-Scr (purple for R1162X-16HBEge, light blue for W1282X-16HBEge) and Ad-Arg (red) or Leu (blue) at MOIs of 10, 50, 100 and 500 and treated with SMG1i (0.3 μM) and/or CFTR modulators as indicated. WT (+ hCAR) transduced with Ad-Scr (MOI 100) served as controls for all experiments (black). (**A, C**) Relative CFTR mRNA abundance of cells collected after Ussing chamber experiment (*n* = 3-6). (**C, F**) Total AUC quantification (*n* = 3-6) of Ussing chamber recordings of CFTR channel function. Data are presented as the mean ± SEM. Breakout analysis for effect size of each treatment are shown in **Figure S4B-S4E**.Significance was determined by one-way ANOVA and Tukey’s post-hoc test, where * *p* < 0.05, ** *p* < 0.01, *** *p* < 0.001 and **** *p* < 0.0001 vs. Ad-Scr MOI 10, (**A, B**) ^##^ *p* < 0.01 and ^####^ *p* < 0.0001 vs. Ad-Arg MOI 10 + LI, and (**C, D**) ^##^ *p* < 0.01 and ^###^ *p* < 0.001 vs. Ad-Leu MOI 10 + ETI.

### ACE-tRNA^Leu^ and ACE-tRNA^Arg^ rescued CFTR channel function with the most prevalent CF-causing nonsense mutations

Next, we sought out to determine if a single ACE-tRNA sequence could be utilized for multiple CF-causing nonsense mutations. Indeed, if one ACE-tRNA isotype can rescue CFTR function for various mutations, clinical application of ACE-tRNAs would be more practical. To this end, Ad-Gly, Ad-Arg and Ad-Leu (MOI 100) were each used to transduce G542X-, R553X-, R1162X- and W1282X-16HBEge cell lines on Transwells, and recovery of CFTR transcript abundance and channel function was subsequently measured. Ad-Leu recovered the most CFTR transcript abundance across all four PTCs (Figure 5A, blue open bars), recovering ∼76% of G542X-, ∼54% of R553X-, ∼60% of R1162X- and ∼59% of W1282X-CFTR, compared to WT 16HBE14o-cells transduced with Ad-Scr (Figure 5A, black open bar). Ad-Arg transduction (Figure 5A, red open bars) recovered the second most transcript abundance across all PTCs with ∼52% of G542X-, ∼49% of R553X-, ∼45% of R1162X- and ∼39% of W1282X-CFTR, compared to WT 16HBE14o-cells transduced with Ad-Scr (Figure 5A, black open bar). For all PTCs, Ad-Gly did not restore transcript abundance significantly (Figure 5A, orange open bars), and as expected ETI treatment did not significantly impact transcript abundance under all conditions (Figure 5A, hashed bars). Interestingly, the rank order of rescued CFTR transcript abundance with Ad-Leu (G542X>R1162X≥W1282X≥R553X) and Ad-Arg (G542X>R553X≥R1162X>W1282X) did not match that of basal transcript expression of CFTR-PTC variants transduced with Ad-Scr (R553X (∼34%) > G542X (∼22%) ≥ W1282X (∼21%) ≥ R1162X (∼17%); Figure 5A), indicating that basal transcript abundance, of which unstimulated PTC readthrough contributes, does not predict suppression efficiency of PTCs within a given transcript.

**Figure 5.**
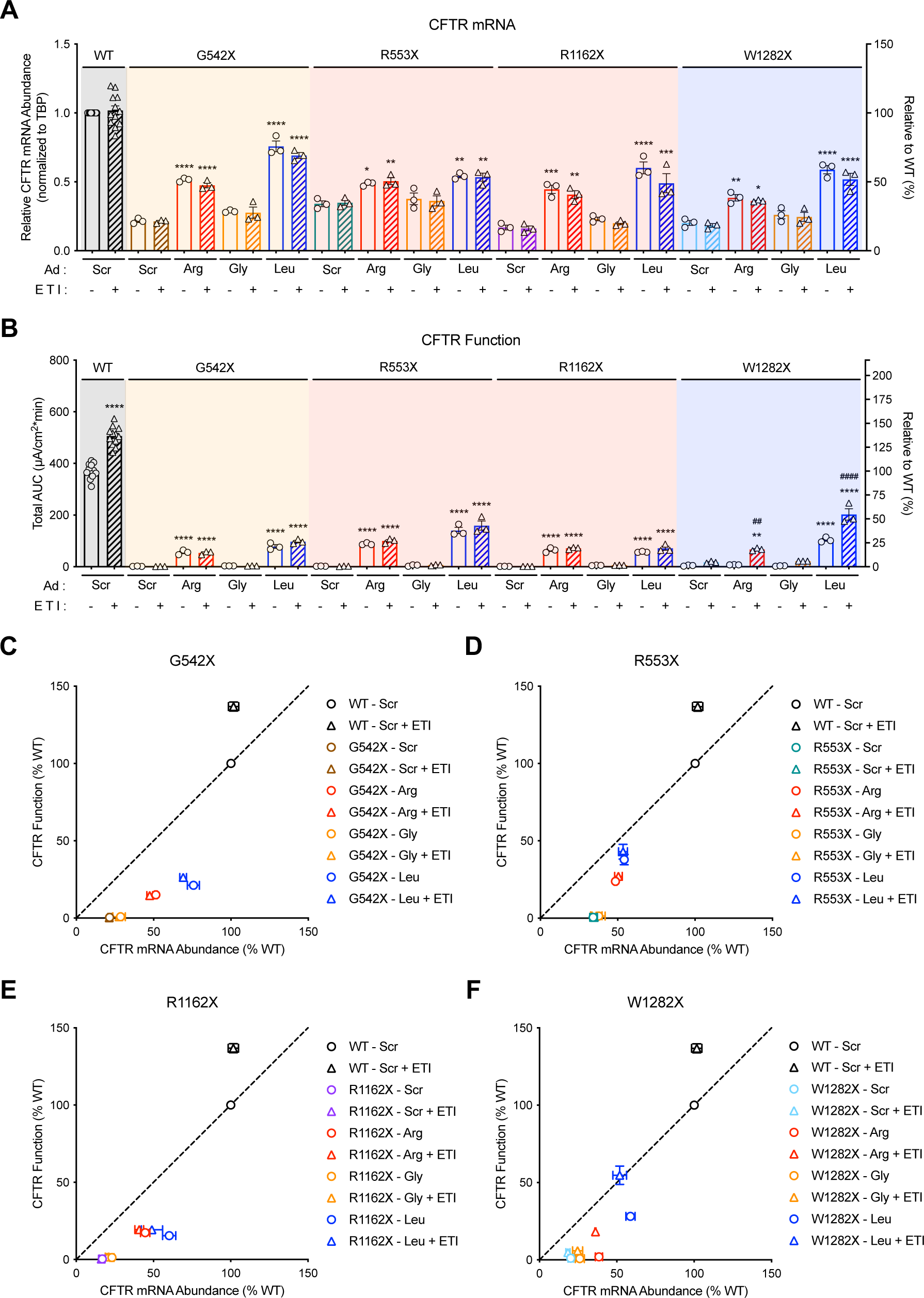
ACE-tRNA^Leu^ significantly rescues CFTR transcript abundance and channel function for the most common CF-causing nonsense variants. (**A-B**) WT, G542X-, R553X-, R1162X-, and W1282X-16HBEge cells (+ hCAR) transduced with Ad-Scr, Ad-Arg (red), Ad-Gly (orange) or Ad-Leu (blue) at an MOI of 100 and treated with either vehicle or ETI. (**A**) Relative CFTR mRNA abundance was quantified after Ussing chamber recordings (*n* = 3-12). (**B**) Total AUC quantification (*n* = 3-12) from Ussing chamber recordings. (**C-F**) CFTR channel function in relation to CFTR mRNA abundance for (**C**) G542X, (**D**) R553X, (**E**) R1162X and (**F**) W1282X cell lines (*n* = 3-12). Breakout analysis for effect size of each treatment on Ussing recordings in (**B**) are shown in **Figure S5A and S5B**. Data are presented as the mean ± SEM. Significance was determined by one-way ANOVA and Tukey’s post-hoc test, where * *p* < 0.05, ** *p* < 0.01, *** *p* < 0.001 and **** *p* < 0.0001 vs. Ad-Scr and ^##^ *p* < 0.01, ^####^ *p* < 0.0001 vs. no ETI.

Despite the generation of leucine substitution mutant CFTR channels in all cell types, Ad-Leu (MOI 100) supported significant rescue of CFTR channel function across all four CFTR PTC 16HBEge cell lines, with ∼21% of WT for G542X-, ∼38% for R553X-, ∼16% for R1162X- and ∼28% for W1282X-CFTR (Figure 5B, open blue bars). Ad-Arg rescued ∼15% of WT for G542X-, ∼24% for R553X- and ∼17% for R1162X-CFTR but did not recover appreciable channel activity in W1282X-16HBEge cells (Figure 5B, open red bars). Interestingly, in all PTC positions except W1282X, the insertion of a leucine or arginine supported marked CFTR channel function. Further, generation of R553L- and R1162L-CFTR variants by Ad-Leu recovered equal or greater CFTR channel function than generating WT CFTR channels with Ad-Arg transduction. Indeed, the CFTR2 database indicates that R1162L is classified as a non-CF causing variant^7^. Using the ACE-tRNA-dependent misincorporation approach, we can now add G542R, G542L, R553L and W1282L as non-CF causing variants. W1282R is classified as a CF-causing missense mutation that is FDA approved for ETI treatment. We therefore tested all arginine (Figure 5B, hashed red bars) and leucine (Figure 5B, hashed blue bars) CFTR variants for ETI dependent functional modulation. In W1282X-16HBEge cells, Ad-Arg recovery increased from ∼2% of WT to ∼18% with the addition of ETI, as predicted by previous ETI theratyping studies^42^. As before, we observed significant enhancement of W1282L-CFTR channel function with ETI, however all other rescued CFTR function were not markedly enhanced by ETI treatment.

When rescued CFTR channel function was plotted against mRNA abundance for each CFTR PTC mutations, it was more apparent that CFTR functional recovery loosely tracked with CFTR transcript abundance (Figures 5C-5F). Outside of Ad-Arg rescue of W1282X, which required ETI treatment for functional rescue, increased CFTR transcript abundance rescue resulted in more functional CFTR channel rescue. However, this relationship was overall shallow for all CFTR PTCs, with more transcript rescued than channel function at all PTC positions with both Ad-Arg and Ad-Leu. Indeed, the only “one-to-one” relationship observed was W1282X-CFTR rescue with Ad-Leu and ETI treatment (Figure 5F, blue triangle). Further, it is unclear with ETI treatment why WT CFTR function was potently enhanced in WT 16HBE14o-cells (Figures 5C-5F, black circles and triangles), while Ad-Arg rescue of R553X- and R1162X-CFTR function was not further enhanced (Figures 5D and 5E, red circles and triangles). Of note, the degree of recovered CFTR channel function in each cell line was near or even surpassed the Ad transduction efficiency for 16HBE cell lines seeded on Transwells (20-30%, Figures S1O-S1V), demonstrating that delivery efficiency is a key limitation to the ACE-tRNA approach.

### Delivery of ACE-tRNA^Leu^ resulted in significant functional rescue of CFTR channels in patient-derived intestinal organoids

Next, we sought to verify ACE-tRNA-dependent PTC suppression efficacy in a primary cell setting by using patient-derived intestinal organoids (PDIOs). First, we optimized the transduction and read-out pipeline, which is summarized in Figure S6A. Of note, Ad vectors utilized in these studies expressed mCherry fluorescent protein instead of GFP. As characterized by mCherry quantification via flow cytometry, transduction of PDIO derived single cells with Ad-Leu resulted in ∼40% transduction efficiency for both MOI 50 and 100 (Figures 6A and S6B). Prior to functional measurements, we measured CFTR transcript abundance to investigate inhibition of NMD by ACE-tRNA^Leu^ suppression activity. In W1282X/W1282X CFTR PDIOs, CFTR transcript abundance increased ∼2.5 fold upon Ad-Leu transduction (MOI 100), which was half of the effect obtained with SMG1i treatment (Figure 6B).

**Figure 6.**
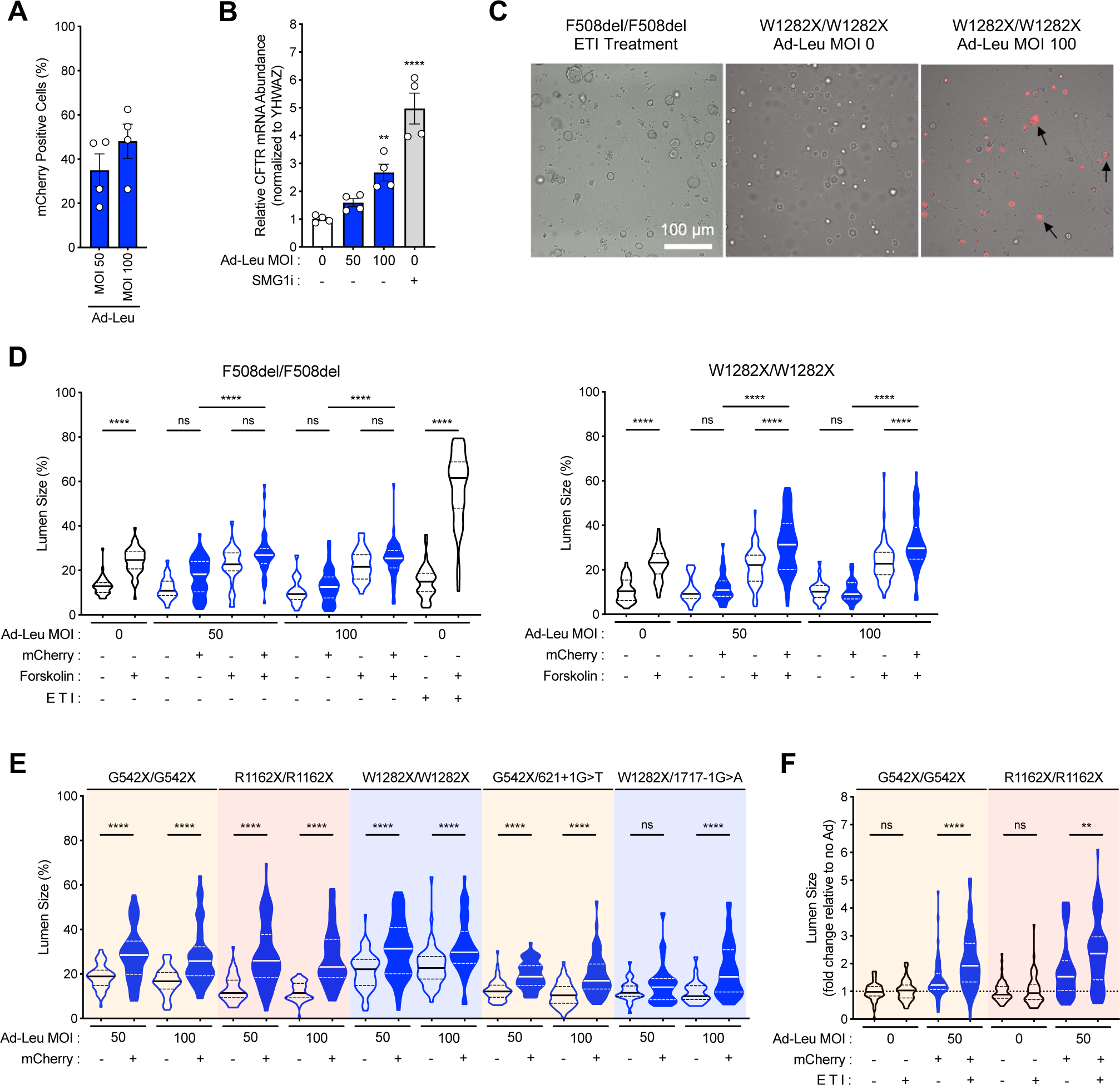
ACE-tRNA^Leu^ significantly recovered CFTR function in patient-derived intestinal organoids (PDIOs) harboring CF-causing nonsense mutations. (**A**) Quantification of mCherry positive cell populations for F508del/F508del and G542X/G542X PIDO-derived single cells transduced with mCherry expressing Ad-Leu at MOIs of 50 and 100. (**B**) Relative CFTR mRNA abundance of W1282X/W1282X PDIOs at 6 DPT upon Ad-Leu transduction or SMG1i (0.3 μM) treatment. (**C**) Representative images of ETI-treated F508del/F508del (left, positive control), untransduced W1282X/W1282X (middle, negative control), and Ad-Leu transduced W1282X/W1282X (right) PDIOs at 6 DPT. Forskolin (5 μM) stimulation at 3 DPT for CFTR activation. Black arrows representing Ad-Leu transduced W1282X/W1282X PDIOs with cystic structures, indicative of the presence of functional CFTR channel. Scale bar, 100 μm. (**D**) Lumen size measurements for F508del/F508del (left, *n* = 22-50) and W1282X/W1282X (right, *n* = 20-60) PDIOs upon transduction with Ad-Leu at MOIs of 50 and 100 in the absence and presence of forskolin stimulation. Ad-Leu transduced PDIOs separated into mCherry negative (Ad-ACE-tRNA untransduced, open violin plot) and mCherry positive (ACE-tRNA transduced, blue filled violin plot) organoid populations. (**E**) Lumen size measurements for five different PDIO genotypes [G542X/G542X (*n* = 38-60), R1162X/R1162X (*n* = 54-70), W1282X/W1282X (*n* = 40-60), G542X/621+1G>T (*n* = 51-61) and W1282X/1717-1G>A (*n* = 26-54)] upon transduction with Ad-Leu at MOIs of 50 and 100. Forskolin stimulation in all conditions for CFTR activation. (**F**) Lumen size relative to Ad untransduced, ETI-untreated condition for G542X/G542X (orange background, *n* = 54-59) and R1162X/R1162X (red background, *n* = 59-60) PDIOs upon transduction with Ad-Leu (MOI 50) in the absence and presence of ETI treatment. Forskolin stimulation in all conditions for CFTR activation. Data are presented as (**A, B**) the mean ± SEM or (**D-F**) violin plots. Significance was determined by one-way ANOVA and (**B**) Dunnett’s post-hoc test, where ** *p* < 0.01 and **** *p* < 0.0001 vs. Ad-Leu MOI 0, or (**D-F**) Tukey’s post-hoc test, where ** *p* < 0.01 and **** *p* < 0.0001.

To allow the outgrowth of single cells into multicellular structures, CFTR was activated with forskolin at 3 days post transduction (DPT) and CFTR function was quantified by measuring lumen area of structures at 6 DPT^43^. As shown in Figure 6C, Ad-Leu resulted in a significant increase in lumen area of W1282X/W1282X CFTR PDIOs upon forskolin stimulation in a subset of the mCherry positive structures. Indeed, when quantifying lumen area in all conditions for W1282X/W1282X CFTR PDIOs, a significant increase in lumen area was present in mCherry positive structures with both MOI 50 and 100 upon stimulation with forskolin (Figure 6D, right). Importantly, in F508del/F508del CFTR PDIOs, which should serve as a negative control, Ad-Leu transduction did not result in a significant increase of luminal size in mCherry positive structures (Figure 6D, left). As expected, ETI treatment of F508del/F508del CFTR PDIOs resulted in a significant increase of lumen area upon forskolin stimulation (Figure 6D, left). To further characterize efficacy of ACE-tRNA^Leu^ in restoring CFTR function, we characterized the efficacy for two additional homozygous PTC genotypes, G542X/G542X and R1162X/R1162X, as well as two compound heterozygous PDIO lines, W1282X/1717-1G>A and G542X/621+1G>T, carrying alleles with CFTR splice mutations that should not benefit from ACE-tRNA activity. Increase in lumen area was significant in mCherry positive structures with an Ad-Leu MOIs 50 and 100 upon stimulation with forskolin and to a similar extent across all three homozygous PTC lines (Figure 6E). However, the increase in lumen area of W1282X/1717-1G>A only reached significance with an Ad-Leu MOI 100 (Figure 6E). Similar to W1282X/W1282X CFTR PDIOs, an increase in lumen area was forskolin dependent (Figure S6C).

We next wanted to determine if ETI treatment could enhance G542L-, and R1162L-CFTR channel function expressed in PDIO cells. In contrast to experiments performed in G542X- and R1162X-16HBEge cells, ETI treatment significantly enhanced Ad-Leu dependent lumen swelling of G542X/G542X CFTR and R1162X/R1162X CFTR PDIO genotypes, while ETI treatment alone did not result in appreciable swelling (Figure 6F). Lastly, we assessed the relation between MFI per organoid structure versus lumen size. We noticed across all experiments that some very bright mCherry positive structures in fact did not show an increased lumen, which could be contributed to Ad vector-associated toxicity upon a high level of transduction. When excluding those structures with the MFI >60 (indicated in gray), a positive linear correlation indicated that higher Ad-Leu transduction resulted in increased PDIO lumen swelling (R=0.28, p<0.0001; Figure S6D).

### ACE-tRNA^Arg^ and ACE-tRNA^Leu^ significantly recovered CFTR channel function in CF patient-derived human primary enteric monolayers

We next determined the efficacy of ACE-tRNA^Arg^ and ACE-tRNA^Leu^ at rescuing CFTR channel function in patient-derived human primary enteric monolayers (hPEMs) cultured on Transwells. hPEMs were first established by Dr. Beekman’s group to study the impact of mutations on CFTR channel function^44^. However, to our knowledge they have not previously been used to study efficacy of CF therapeutic approaches. As observed by representative I_eq_ traces measuring CFTR function, transduction of compound heterozygous G542X/R1162X CFTR hPEMs with increasing MOI (100 to 400) of Ad-Arg (red lines) and Ad-Leu (blue lines) resulted in a saturating and significant rescue of CFTR channel function (Figure 7A). Importantly, there was no channel activity observed with vehicle + I and Ad-Scr MOI 600 + I treatments (Figure 7A, black traces). Based on the average responses of three technical replicates plotted in Figure 7B, Ad-Arg (red bars) and Ad-Leu (blue bars) at higher MOIs rescued similar CFTR channel function as G418 treatment (100μM, green bars). In hPEMs with G542X/R553X (Figure 7C) and W1282X/W1282X (Figure 7D) CFTR genotypes, Ad-Leu (blue bars) rescued significant CFTR channel function at MOIs 300 and 400, however only 30-40% of functional rescue achieved by G418 treatment (green bars). Ad-Arg rescued significant CFTR function in G542X/R553X CFTR hPEMs (Figure 7C, red bars), but not in W1282X/W1282X hPEMs (Figure 7D, red bars), as expected with the given CFTR genotypes. Unexpectedly, for all genotypes, Ad-Arg and Ad-Leu rescued CFTR channel function was not significantly enhanced by ETI treatment (Figure 7B-7D, hashed bars). The representative *I*_eq_ traces for results in Figures 7C and 7D are shown in Figures S7A and S7B.

**Figure 7.**
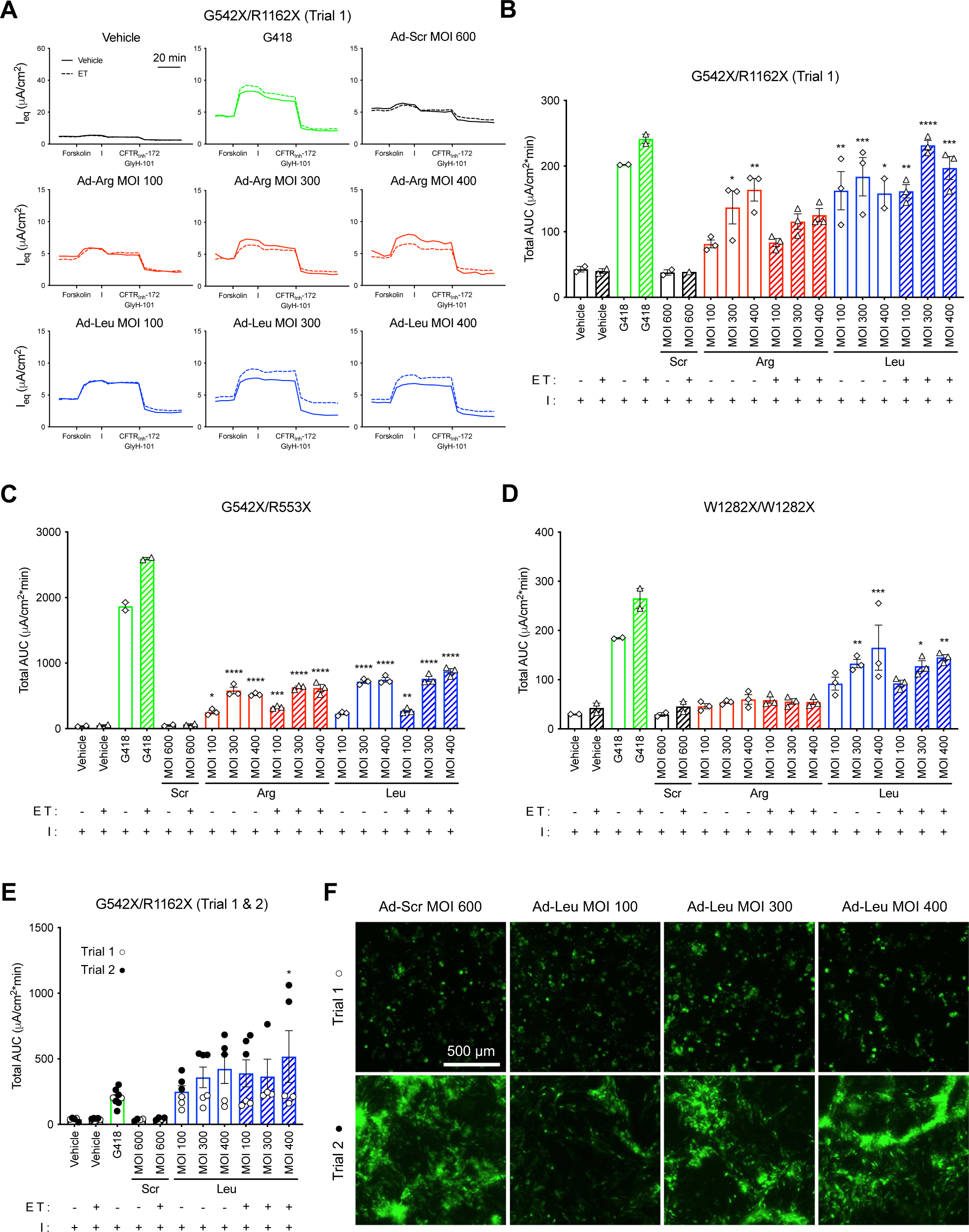
ACE-tRNA^Arg^ and ACE-tRNA^Leu^ significantly recovered CFTR channel function in CF patient-derived human primary enteric monolayers (hPEMs). hPEMs transduced with Ad-Scr (black), Ad-Arg (red) or Ad-Leu (blue) or treated with G418 (100 μM), and subsequently ETI. (**A**) Representative *I*_eq_ traces and (**B**) total AUC quantification (*n* = 2-3) of G542X/R1162X hPEMs (Trial 1). *I*_eq_ traces from TECC-24 assays in response to sequential addition of forskolin (10 μM) and IBMX (100 μM), followed by ivacaftor (I, 1 μM), and then inhibited by a combination of CFTR_Inh_-172 (20 μM) and GlyH-101 (20 μM). Total AUC quantifications of (**C**) G542X/R553X (*n* = 2-3) or (**D**) W1282X/W1282X (*n* = 2-3) hPEMs. (**E**) Total AUC quantification for Combined Trial 1 (open circles) and Trial 2 (closed circles) (*n* = 2-3/trial) of G542X/R1162X hPEMs. (**F**) Representative GFP fluorescence images of Ad-transduced G542X/R1162X hPEMs from first (top) and second (bottom) trials at 3 DPT. (**B-D**), data are presented as the mean ± SEM. Significance was determined by one-way ANOVA and Tukey’s post-hoc test, where * *p* < 0.05, ** *p* < 0.01, *** *p* < 0.001 and **** *p* < 0.0001 vs. Ad-Scr MOI 100 + I.

Given hPEMs are a new culture model for studying gene therapy approaches, we were interested in the overall variability of CFTR functional rescue between Transwell seedings. We therefore re-seeded G542X/R1162X CFTR hPEMs on Transwells, and repeated both G418 treatment and Ad-Leu transduction studies. In “trial 2” (black dots), the magnitude of CFTR channel rescue was significantly improved compared to “trial 1” at all Ad-Leu MOIs tested (Figure 7E, white dots) and could be significantly enhanced by ETI treatment (Figure 7E, black dots over hashed blue bars), while the G418-dependent rescue was similar between trials. The average Ad-Leu dependent rescue of CFTR function between “trial 1” and “trial 2” are represented by the blue bars (I treatment, open bars; ETI treatment, hashed bars). To determine if overall transduction efficiency contributed to the disparity in functional rescue between trials, we performed fluorescent imaging of the Ad GFP expression marker in the hPEMs in the same cells following electrophysiology measurements. Indeed, there was a large disparity in transduction efficiency between trials, with significantly more GFP fluorescence in “trial 2” compared to “trial 1” (Figure 7F). Thus, increased Ad-Leu transduction efficiency was likely the reason for increased rescue of G542X/R1162X CFTR channel function in “trial 2”. Indeed, hPEMs are a promising cell model for studying nonsense CF therapeutics, but do carry the well documented limitation observed in nearly all primary cell models, with variability in phenotypes across cell donors or even within technical replicates^45^. Our study highlights the value of repeated technical replicates and appropriate controls when using primary cell models.

Overall, this study demonstrates the discovery of broad-spectrum ACE-tRNA for multiple nonsense mutation sites with different PTC contexts as well as native amino acids and the ability of modulator drugs in assisting the functional protein recovery. In combination with an efficient delivery system, ACE-tRNA technology may be a promising therapeutic option for genetic disease caused by nonsense mutations in a CF-overarching manner.

## DISCUSSION

There are considerable ongoing efforts to develop therapeutics that target disease causing PTCs as a PTC-agnostic technology would have the potential to treat millions of individuals with serious PTC-associated diseases. As a PTC-agnostic therapeutic avenue, aminoglycosides and other PTC readthrough inducing small molecules have been primarily studied. Their general ability to induce readthrough at all three PTCs (UGA, UAG and UAA) and generate full-length protein is attractive, however they either have shown toxicity through off-target mechanisms or show low efficacy *in vivo*^13^. We recently generated a library of ACE-tRNAs that allows for the rescue of WT protein sequence and function from all disease causing nonsense mutations^21^. Here, we shifted our focus from generating ‘seamless’ rescued full-length proteins to a more PTC agnostic approach, akin to small molecule readthrough agents. Specifically, we harnessed the ability of ACE-tRNAs to precisely suppress one of three PTCs with a specific amino acid to determine if a single ACE-tRNA isotype could be utilized for several CF-causing nonsense mutations. We found that our best-performing ACE-tRNA that decodes all UGA PTCs to the leucine amino acid (ACE-tRNA^Leu^) resulted in significant recovery of endogenous CFTR channel function from the most common CF nonsense mutations. These findings support ACE-tRNAs as a platform nonsense therapeutic for rescue of protein function encoded from any gene with PTCs arising from multiple different parent codon identities. We highlighted in this study with W1282X-CFTR that suppression of a PTC with the ‘incorrect’ amino acid, in this case arginine, resulted in a non-functional CFTR channel (Figure 5B). Knowingly, we generated a CF missense mutation W1282R-CFTR that could be rescued with subsequent ETI treatment. Indeed, these findings highlight the benefit of an ACE-tRNA-modulator combinatorial therapeutic approach for CF. Interestingly, addition of NMD inhibitor to the ACE-tRNA CFTR modulator combination increased CFTR mRNA level but minimally enhanced functional channel level, indicating that ACE-tRNAs are very efficient at PTC suppression and the presence of additional transcripts due to NMD inhibition has not much additional benefit on CFTR function. We speculate that PTC readthrough efficiency is the same mechanistic pinch point for recovery of transcript abundance during the pioneer round of translation and production of full-length protein from stabilized transcripts. Therefore, increasing PTC-containing transcript abundance through NMD inhibition may not provide significant benefit with combinatorial use with ACE-tRNAs, as it has been observed with less active readthrough agents such as aminoglycosides^27,41^. Regardless, circumventing the need for general NMD inhibition for nonsense therapeutics is highly favorable due to concerns of toxicity^46^.

To determine broad applicability of one ACE-tRNA isotype to rescue functional protein from several genes harboring multiple PTCs, a theratyping approach will need to be used on a PTC-to-PTC basis to determine efficacy of rescue^47^. Likely, there will be proteins that are not as amendable to the incorporation of non-WT amino acids into PTCs, where missense mutations have shown to result in non-functional protein and disease. A disputation to this idea is *CFTR*, where 40% of CF causing mutations are missense^12,48^. However, there are several disorders where missense mutations are found to be uncommon. Two examples are Duchenne muscular dystrophy and titinopathies, where truncating deletions, duplications and nonsense and frameshift mutations in dystrophin (*DMD*) and titin (*TTN*) genes, respectively, are the most common variants^49,50^.

Total gene complementation, that is delivery or replacement of the correct gene product, is considered the ideal genetic therapeutic approach. A promising gene complementation therapeutic approach is mRNA delivery to the target tissue. With mRNA, entry into the cell is the only barrier for expression, making it an attractive option for non-dividing cells. However, the relatively short half-life of mRNA will necessitate frequent redosing^51^. If viral delivery is utilized, the transgene expression cassette must be under the payload limits of therapeutic viruses such as AAV (<5kb) and lentivirus (<8kb). While maneuvers can be made to reduce the size of the therapeutic transgene, as exemplified with CFTR^52^ and DMD^53^, not every protein is amendable to such engineering. ACE-tRNA transgenes are small, with an entire expression cassette size of 150-200bps^23,31^, and can be readily encoded in AAVs^22^. Further, ACE-tRNAs can be delivered as RNA and DNA, opening up delivery options for targeting different tissues^23,54^.

Base editing technologies to revert the genomic PTC to a coding codon and mRNA PTC pseudouridylation to promote co-translational near-cognate tRNA suppression have both been used to effectively rescue full-length proteins from genes harboring PTCs^55–58^. These approaches use the PTC nucleotide context to gain specificity in PTC targeting, thus limiting potential off-target activity. As with ACE-tRNAs, the rescued protein expression remains under endogenous control which is important for genes that require tight physiologic regulation. With these PTC targeted approaches, the compromise is the necessity to develop a tailored therapeutic for each PTC. Sup-tRNA technologies have been used to effectively recover full-length protein expression from several genes of different sizes and function from several disease causing PTCs, including but not limited to *HBB* (β-thalassaemia)^17^, *CFTR* (cystic fibrosis)^21,24,26^, *DMD* (Duchenne muscular dystrophy)^59^, *KCNH2* (long-QT syndrome)^60^, *XPA* (Xeroderma pigmentosum)^61^ and *IDUA* (mucopolysaccharidosis type I)^22^. Here, we demonstrate that ACE-tRNAs can be utilized more broadly without focusing on generating the WT protein for therapeutic effect, therefore being limited by the specific PTC codon, UGA, UAA or UAG. We purposefully utilized ACE-tRNA^Leu^ in this study, as it is one of two ACE-tRNA families (i.e. isoacceptor) identified in our original screen, the other being ACE-tRNA^Ser^ with three different highly-functional ACE-tRNAs specific for each PTC (UGA, UAG and UAA)^21^. Indeed, using ACE-tRNA^Leu^ and ACE-tRNA^Ser^, the platform therapeutic approach demonstrated here can be readily tested with all PTCs in every gene that results in a nonsense-associated diseases. Indeed, it is an exciting prospect to identify a single ACE-tRNA variant for each PTC codon as an effective therapeutic for several, if not most nonsense-associated diseases.

## ACKNOWLEDGMENTS

We thank members of the Lueck Laboratory for editing the manuscript and their constructive discussion throughout the study. We also thank Dr. Amy M. Martin for reading the manuscript for grammatical errors and T.N. Lueck with help generating figures. We would like to thank the Cystic Fibrosis Foundation Therapeutics Lab and Dr. Hillary Valley for providing 16HBE14o- and 16HBE14ge cell lines used in this study. This work was supported by Emily’s Entourage by a grant to S.P, J.M.B. and J.D.L., Cystic Fibrosis Foundation Postdoctoral Fellowship (PORTER20F0) to J.J.P., and a Cystic Fibrosis Foundation Research Grant (LUECK20GO), and NIH grant (1 R01 HL153988) to J.D.L.

## AUTHOR CONTRIBUTIONS

W.K., J.J.P., S.S., T.C., P.B., K.C., J.M.B. and J.D.L. designed the study. W.K., J.J.P., S.S., T.C. P.B., I.W. and J.D.L. performed experiments. W.K., J.J.P., S.S., T.C. P.B., I.W. and J.D.L. analyzed the data and constructed the figures. W.K., J.J.P., S.S. and J.D.L. wrote the manuscript. All authors read and revised the manuscript.

