## Supplemental Figures 1-7 and Legends for "Anticodon Engineered Transfer RNAs (ACE-tRNAs) are a Platform Technology for Suppressing Nonsense Mutations"

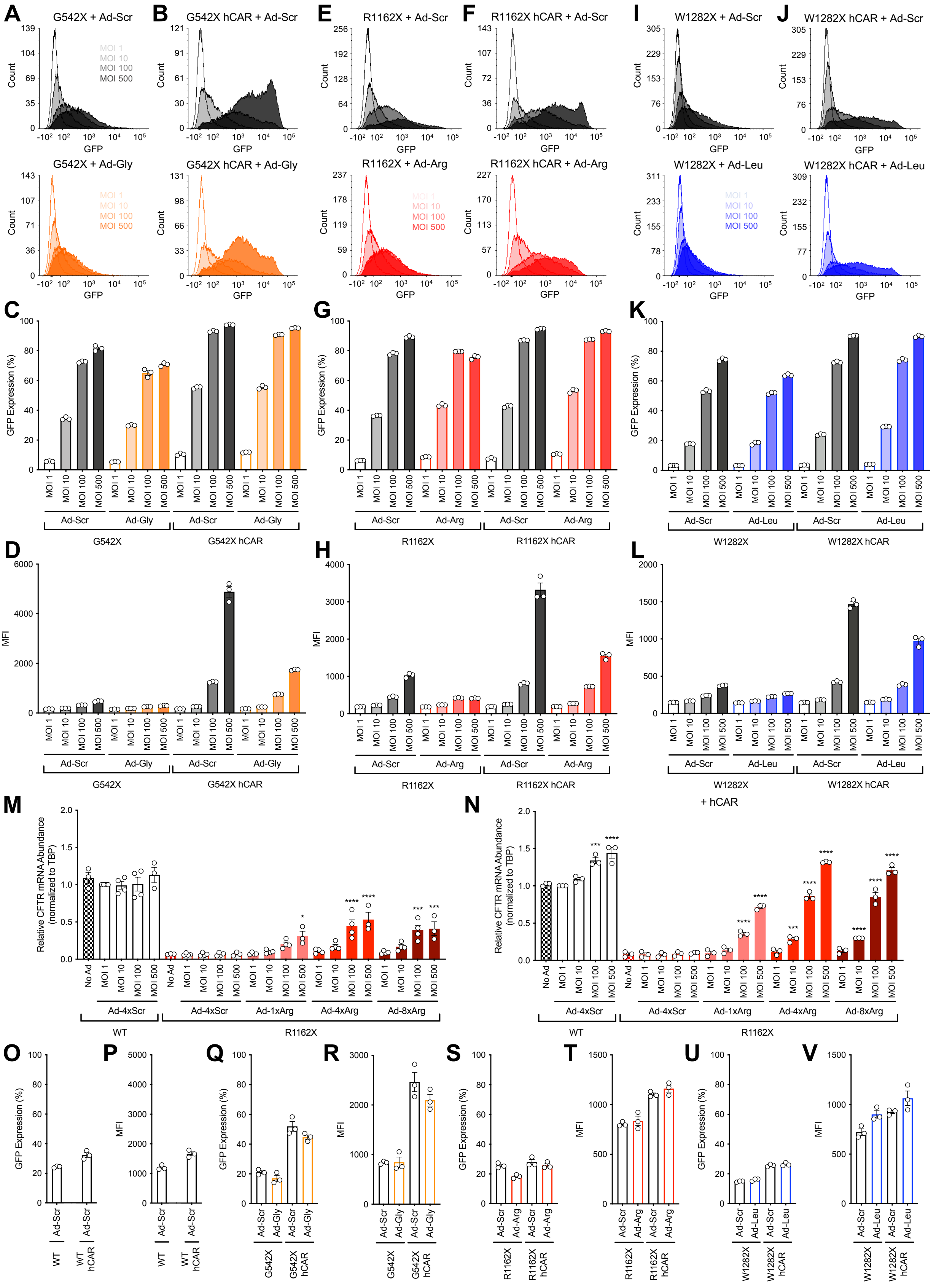


### Supplemental Figure 1

**Figure S1. Characterization of Ad-ACE-tRNA transduction efficiency of HBE cells. Related to Figure 1.** (**A-D**) Representative flow cytometry histograms of (**A**) G542X-16HBEge or (**B**) + hCAR cells transduced with Ad-Scr (top, black) or Ad-Gly (bottom, orange) at different MOIs. Quantification of (**C**) GFP positive cell population and (**D**) median fluorescence intensity (MFI) of that population from flow cytometry data ($n = 3$). (**E-H**) Representative flow cytometry histograms of (**E**) R1162X-16HBEge or (**F**) + hCAR cells transduced with Ad-Scr (top, black) or Ad-Arg (bottom, red) at different MOIs. Quantification of (**G**) GFP positive cell population and (**H**) MFI of that population from flow cytometry data ($n = 3$). (**I-L**) Representative flow cytometry histograms of (**I**) W1282X-16HBEge or (**J**) + hCAR cells transduced with Ad-Scr (top, black) or Ad-Leu (bottom, blue) at different MOIs. Quantification of (**K**) GFP positive cell population and (**L**) MFI of that population from flow cytometry data ($n = 3$). (**M-N**) Relative CFTR mRNA expression in (**M**) original WT and R1162X-16HBEge cells ($n = 3$-4) or (**N**) + hCAR cells ($n = 3$) transduced with different MOIs of Ad-Scr or Ad-Arg harboring different copy number of the ACE-tRNA$^{Arg}$ expression cassette (1x for 1 copy, 4x for 4 copies and 8x for 8 copies). (**O-V**) Quantification of (**O, Q, S, U**) GFP positive cell population and (**P, R, T, V**) MFI from flow cytometry analysis of (**O-P**) WT and WT + hCAR, (**Q-R**) G542X-16HBEge and G542X-16HBEge + hCAR, (**S-T**) R1162X-16HBEge and R1162X-16HBEge + hCAR and (**U-V**) W1282X-16HBEge and W1282X-16HBEge hCAR cells plated on Transwells and transduced with Ad-Scr or Ad-ACE-tRNAs as indicated in the figure at MOI of 100 ($n = 3$). Data are presented as the mean ± SEM. For (**M, N**), significance was determined by one-way ANOVA and Tukey's post-hoc test, where *$p < 0.05$, ***$p < 0.001$ and ****$p < 0.0001$. vs. Ad-4xScr MOI 1.

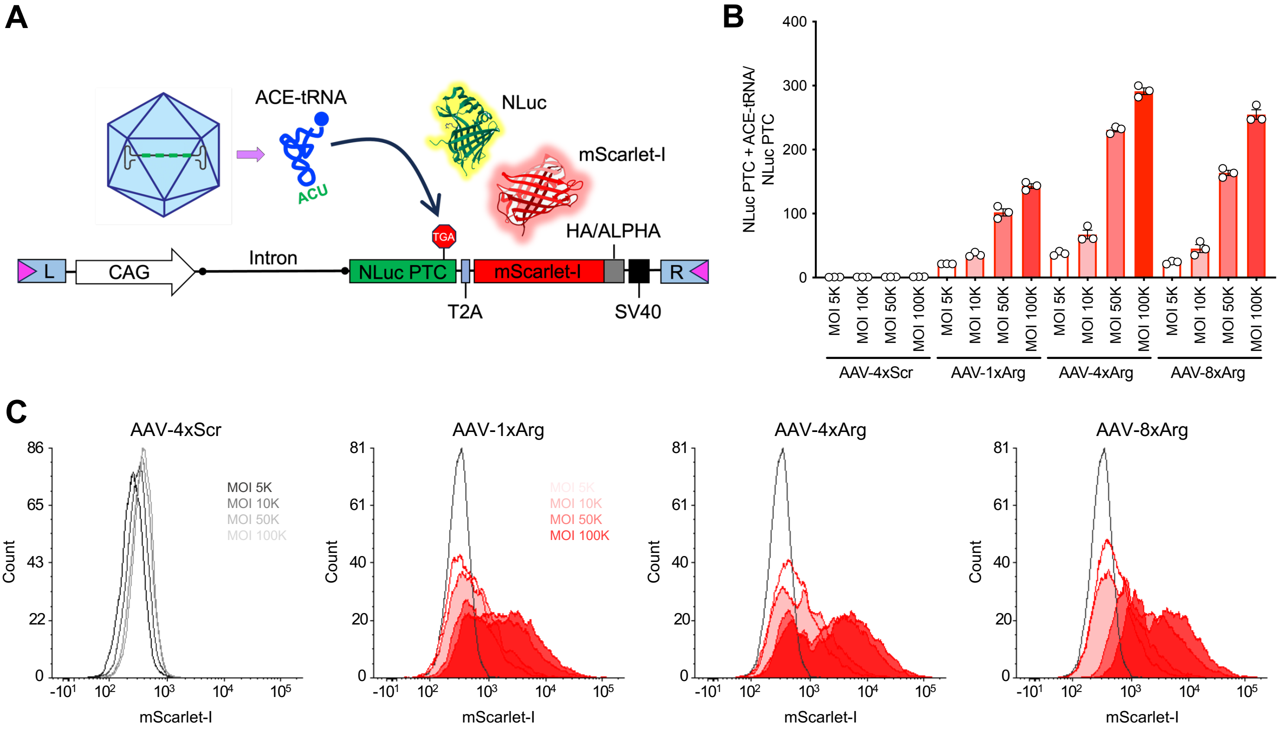


### Supplemental Figure 2

**Figure S2. Efficient delivery of ACE-tRNAs by AAV2.** (**A**) Schematic of the *piggyBac* stop-go-glow (SGG) PTC reporter cassette with an in-frame PTC-containing NanoLuc (NLuc) luciferase and a mScarlet-I fluorescent protein. (**B**) Normalized luminescence and (**C**) representative flow cytometry histograms of the SGG PTC reporter stably integrated HEK293T cells transduced with AAV2-Scr (black) or AAV2-Arg (red) at different MOIs.

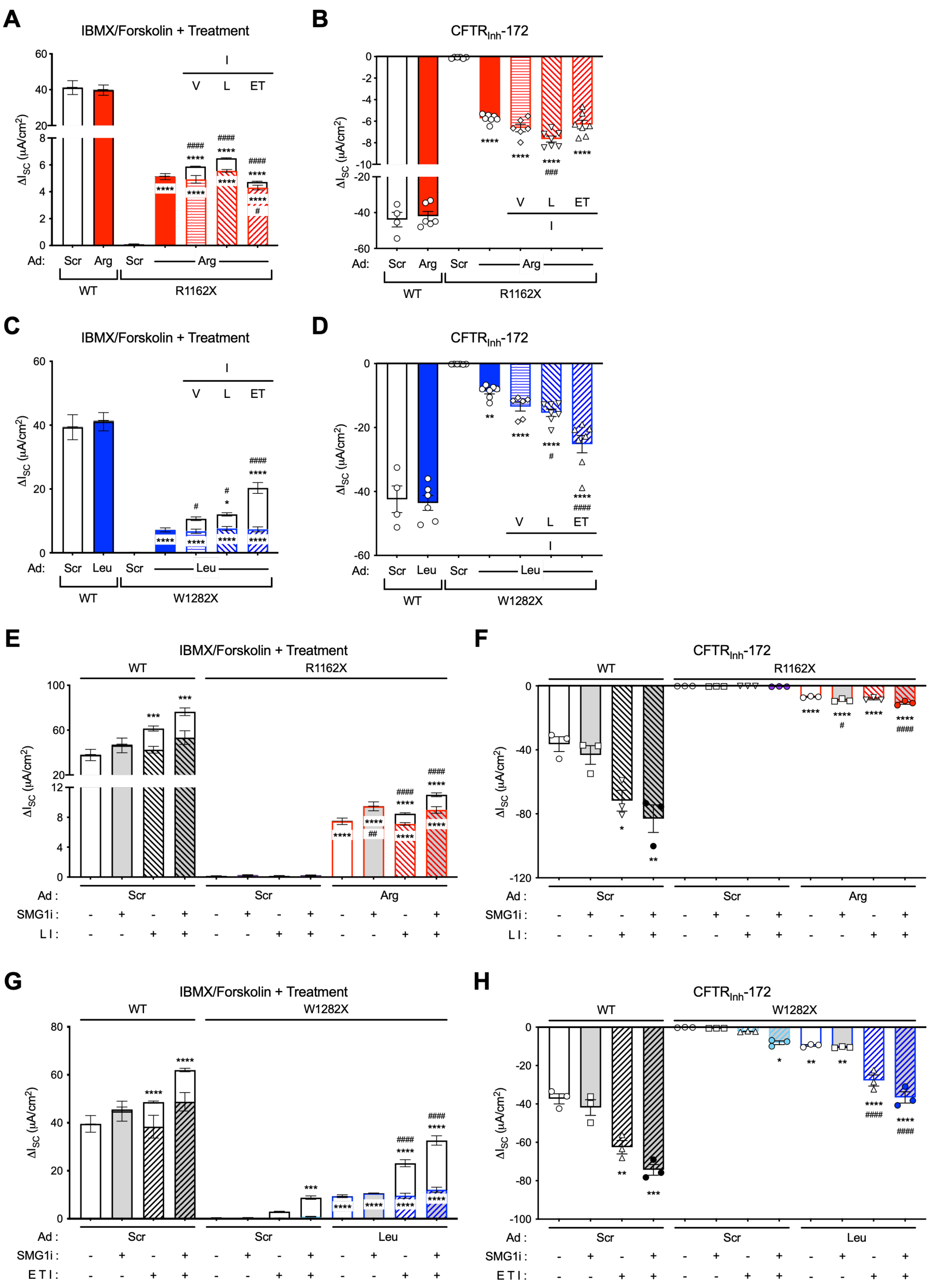


### Supplemental Figure 3

**Figure S3. CFTR modulators potentiate ACE-tRNA recovered CFTR function. Related to Figures 2 and 3.** (**A-B, E-F**) R1162X-16HBEge and (**C-D, G-H**) W1282X-16HBEge cells (+ hCAR) transduced with Ad-Scr and Ad-Arg (red) or Leu (blue) at an MOI of 100 and treated with indicated CFTR modulators [vehicle (V), lumacaftor (L), elexacaftor and tezaxaftor (ET), Ivacaftor (I)]. Gray filled bar representing SMG1i (0.3 $\mu$M) treated conditions for (**E-H**). WT (+ hCAR) served as control for all experiments. Maximum $I_{SC}$ change in response to sequential addition of (**A, C, E, G**) forskolin and IBMX (bottom bar), vehicle or ivacaftor (top bar) and (**B, D, F, H**) $CFTR_{Inh}$-172 for the experiment presented in **Figures 2B, 2E, 3D and 3H** ($n$ = 3-7). Data are presented as the mean ± SEM. Significance was determined by unpaired t-test or one-way ANOVA and Tukey's post-hoc test, where $^{*}p < 0.05$, $^{**}p < 0.01$, $^{***}p < 0.001$ and $^{****}p < 0.0001$ vs. Ad-Scr, $^{\#}p < 0.05$, $^{\#\#}p < 0.01$, $^{\#\#\#}p < 0.001$ and $^{\#\#\#\#}p < 0.0001$ vs. Ad-Arg, and $^{\#}p < 0.05$ and $^{\#\#\#\#}p < 0.0001$ vs. Ad-Leu.

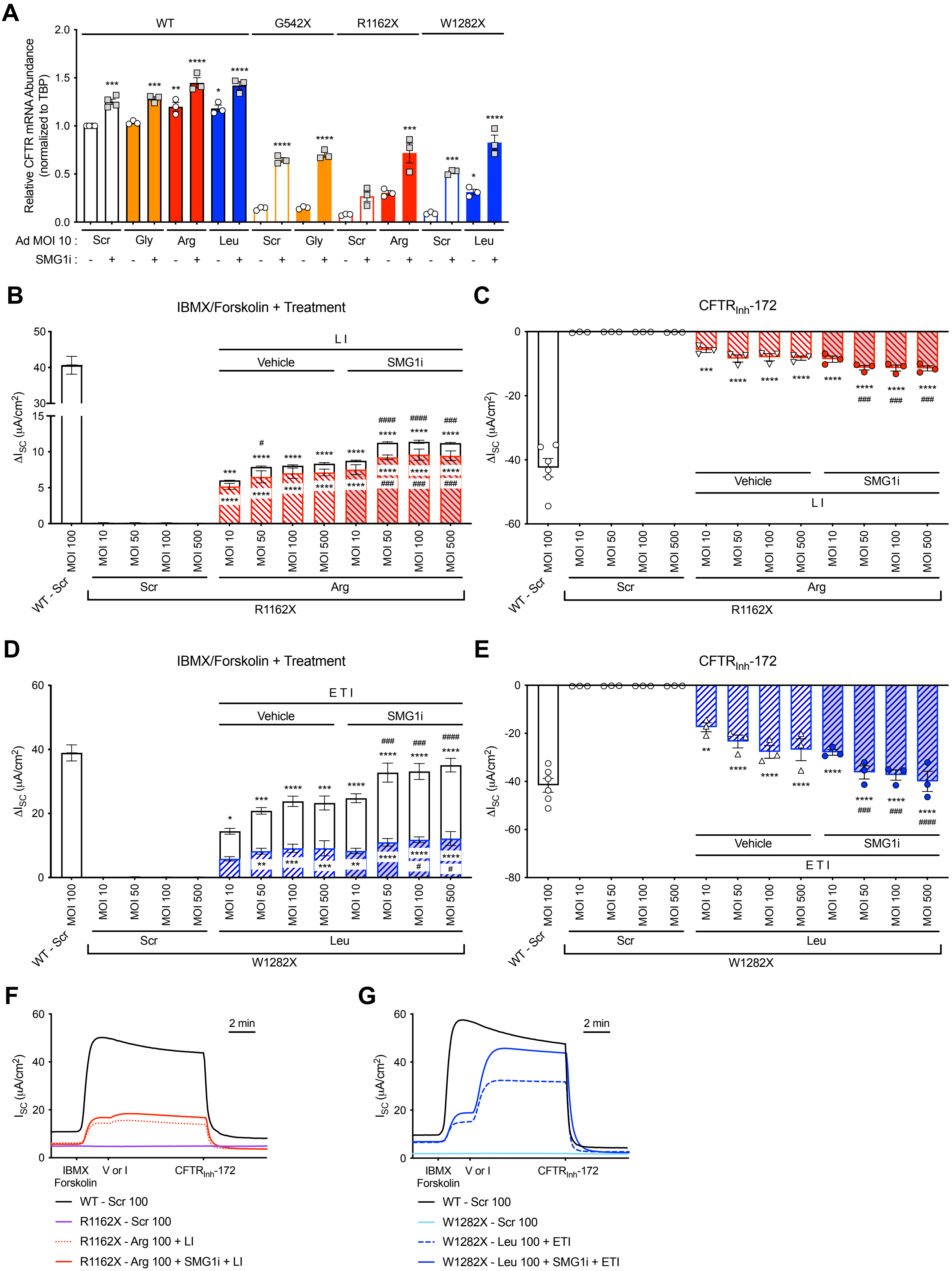


### Supplemental Figure 4

**Figure S4. Combinatorial treatment of ACE-tRNAs with CFTR modulators and NMD inhibition. Related to Figure 4.** (**A**) CFTR mRNA abundance normalized to TBP in WT, G542X-, R1162X-, and W1282X-16HBEge cells (+ hCAR) transduced with Ad-Scr (open bar), Ad-Gly (orange filled bar), Ad-Arg (red filled bar) or Ad-Leu (blue filled bar) at MOI of 10 in the presence (+) and absence (-) of SMG1i (0.3 μM) treatment as indicated in the figure ($n = 3$-4). (**B-C, F**) R1162X16HBEge or (**D-E, G**) W1282X-16HBEge cells (+ hCAR) transduced with Ad-Scr (black for WT, purple for R1162X-16HBEge, light blue for W1282X-16HBEge) and Ad-Arg (red) or Ad-Leu (blue) at MOIs of 10, 50, 100 and 500 and treated with SMG1i and/or CFTR modulators as indicated in the figure. WT (+ hCAR) served as control for all experiments. Maximum $I_{SC}$ change in response to sequential addition of (**B, D**) forskolin and IBMX (bottom bar), vehicle or ivacaftor (top bar) and (**C, E**) $CFTR_{Inh}$-172 for the experiment presented in (**F, G**) representative $I_{SC}$ traces for MOI 100 ($n = 3$-6). Data are presented as the mean ± SEM. Significance was determined by one-way ANOVA and Tukey's post-hoc test, where $^{**}p < 0.05$, $^{**}p < 0.01$, $^{***}p < 0.001$ and $^{****}p < 0.0001$ vs. Ad-Scr MOI 10, $^{\#}\ p < 0.05$, $^{\#\#\#}\ p < 0.001$ and $^{\#\#\#\#}\ p < 0.0001$ vs. Ad-Arg MOI 10 + LI and $^{\#}\ p < 0.05$, $^{\#\#\#}\ p < 0.001$ and $^{\#\#\#\#}\ p < 0.0001$ vs. Ad-Leu MOI 10 + ETI.

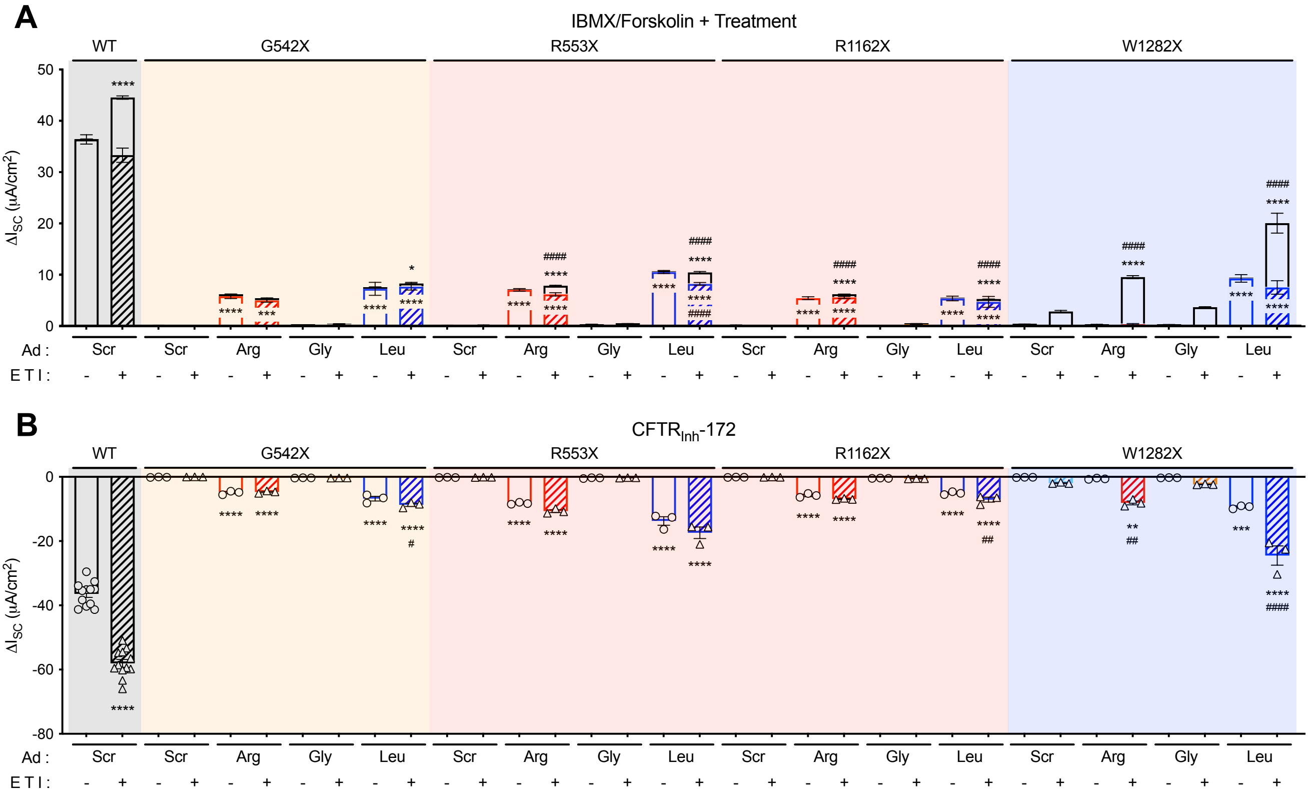


### Supplemental Figure 5

**Figure S5. ACE-tRNA$^{Leu}$ significantly rescues functional CFTR level in the most prevalent CF-causing nonsense mutations. Related to Figure 5.** WT, G542X-, R553X-, R1162X-, and W1282X-16HBEge cells (+ hCAR) transduced with Ad-Scr, Ad-Arg (red), Ad-Gly (orange) or Ad-Leu (blue) at an MOI of 100 and treated with either vehicle or ETI. Maximum $I_{SC}$ change in response to sequential addition of (**A**) forskolin and IBMX (bottom bar), vehicle or ivacaftor (top bar) and (**B**) CFTR$_{Inh}$-172 ($n$ = 3-12). Data are presented as the mean ± SEM. Significance was determined by unpaired t-test or one-way ANOVA and Tukey's post-hoc test, where *$p < 0.05$, **$p < 0.01$, ***$p < 0.001$ and ****$p < 0.0001$ vs. Ad-Scr and $^{\#}$ $p < 0.05$, $^{\#\#}$ $p < 0.01$, $^{\#\#\#\#}$ $p < 0.0001$ vs. no ETI.

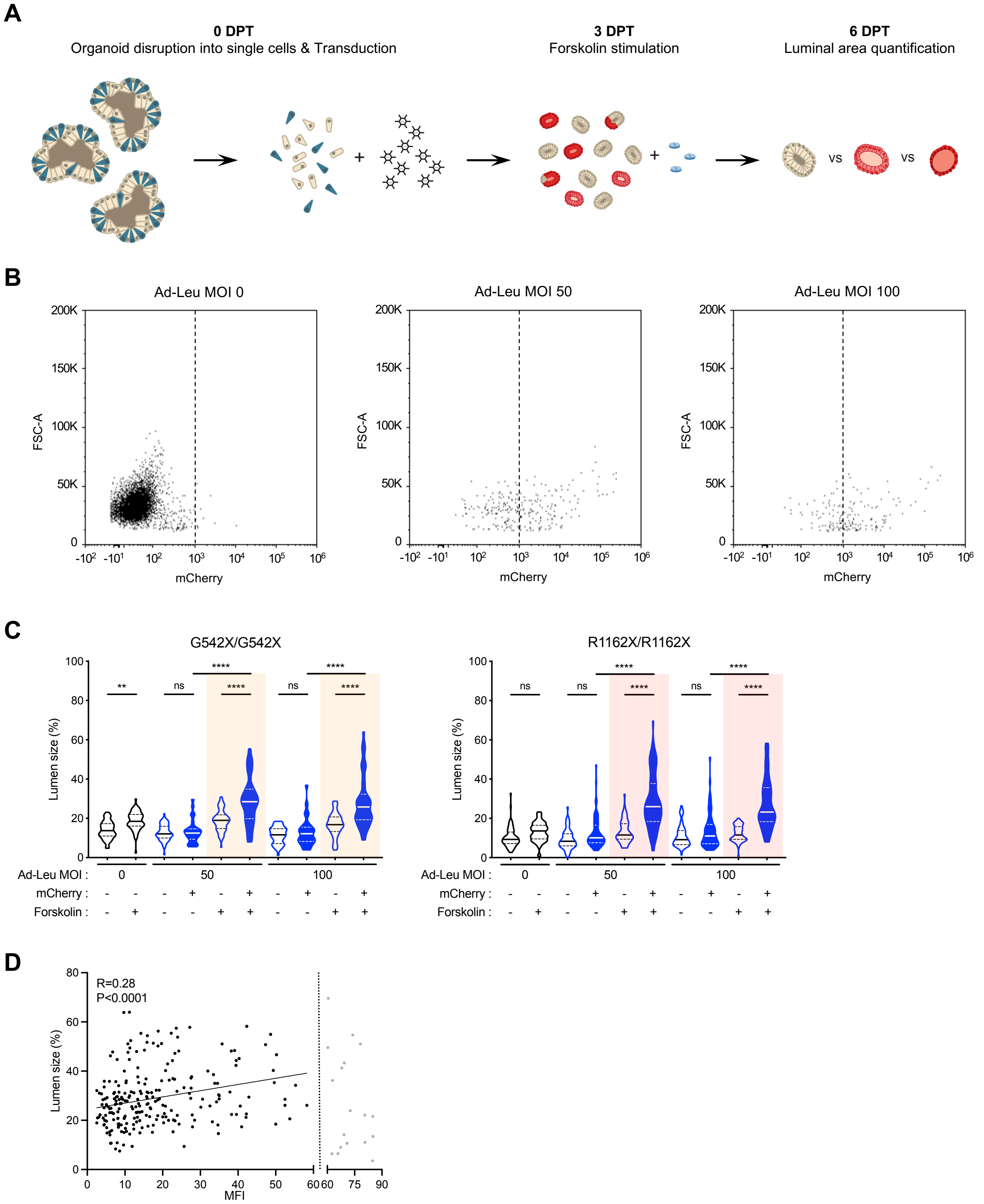


### Supplemental Figure 6

**Figure S6. Ad-ACE-tRNA$^{Leu}$ transduction of PDIOs rescues CFTR channel function as indicated by increased luminal area upon forskolin stimulation. Related to Figure 6.** (**A**) Schematic of PDIO transduction pipeline and experimental timeline. Stem cells indicated in blue and mCherry positive cells indicated in red. (**B**) Representative flow cytometry plots of W1282X/W1282X PDIOs upon Ad-Leu transduction at MOIs of 50 (middle) or 100 (right) compared to untransduced condition (left) at 6 DPT. Dotted line represents the gate for determining mCherry negative (ACE-tRNA untransduced) and positive (ACE-tRNA transduced) organoid populations. (**C**) Lumen size measurements for G542X/G542X (left, $n = 27$-63) and R1162X/R1162X (right, $n = 49$-120) PDIOs upon transduction with Ad-Leu at MOIs of 50 or 100 in the absence and presence of forskolin (5 μM) stimulation. Ad-Leu transduced PDIOs separated into mCherry negative (ACE-tRNA untransduced, open violin plot) and mCherry positive (ACE-tRNA transduced, blue filled violin plot) organoid populations. Data with colored background (part of **Figure 6E**) included for comparison with other conditions. Data are presented as violin plots. Significance was determined by one-way ANOVA and Tukey's post-hoc test, where $^{**}p < 0.01$ and $^{****} p < 0.0001$. (**D**) The correlation between MFI per structure (X-axis) and lumen size (Y-axis) for mCherry positive structures of three homogenous PTC PDIOs (G542X/G542X, R1162X/R1162X and W1282X/W1282X) upon Ad-Leu transduction at MOIs of 50 and 100 and forskolin stimulation. Gray dots indicate organoids with MFI > 60 that were excluded for a linear regression analysis, which results in a positive correlation with R=0.28 and $p < 0.0001$.

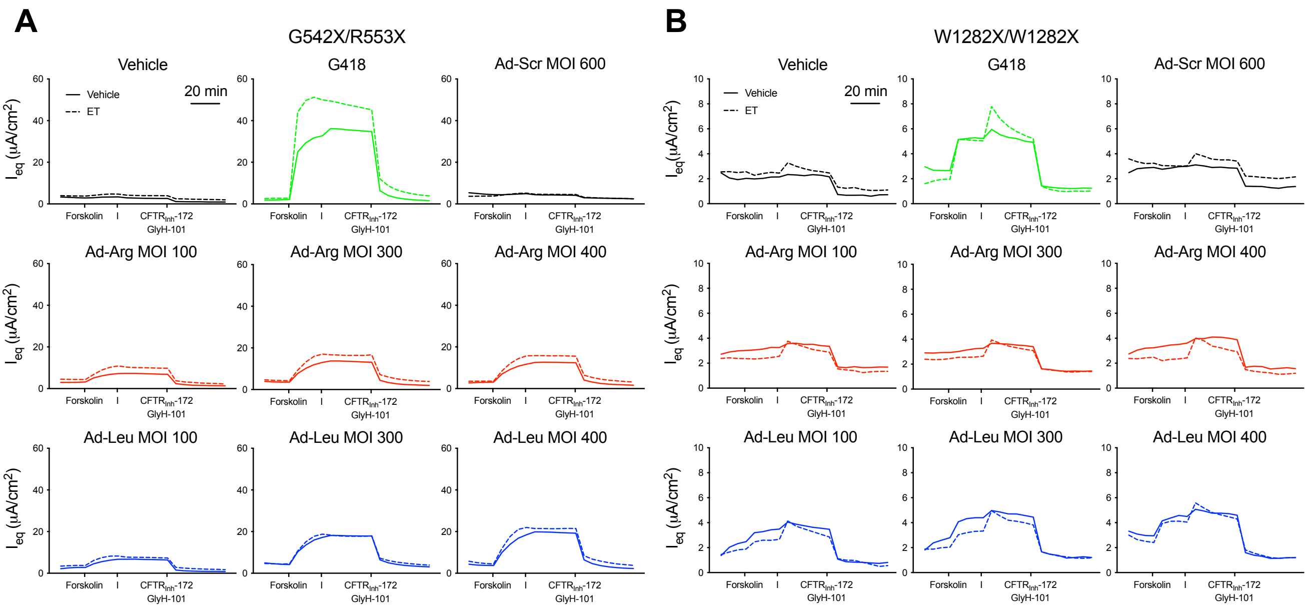


### Supplemental Figure 7

**Figure S7. Ad-ACE-tRNA$^{Arg}$ and Ad-ACE-tRNA$^{Leu}$ transduction of hPEMs recovers CFTR channel function. Related to Figure 7.** Representative $I_{eq}$ traces for the experiment presented in **Figures 7C and 7D** of (**A**) G542X/R553X and (**B**) W1282X/W1282X hPEMs transduced with Ad-Scr (black), Ad-Arg (red) and Ad-Leu (blue) and treated with vehicle (solid line) or CFTR correctors (ET, dashed line). Representative $I_{eq}$ traces in response to sequential addition of forskolin (10 μM) and IBMX (100 μM), followed by ivacaftor (I, 1 μM), and then inhibited by a combination of CFTR$_{Inh}$-172 (20 μM) and GlyH-101 (20 μM).
